# Investigating Age-Related Decline in Sensorimotor Control Using Robotic Tasks

**DOI:** 10.1101/2025.07.07.663471

**Authors:** Laura Alvarez-Hidalgo, Ellie Edlmann, Gunnar Schmidtmann, Ian S. Howard

## Abstract

Aging is associated with changes in sensorimotor control that contribute to functional decline, mobility limitations, and increased fall risk. Traditional motor assessments often rely on subjective measures, highlighting the need for objective, quantitative tools. We developed three robot-based tasks using the vBOT planar manipulandum to evaluate sensorimotor performance in healthy young (<35 years) and older (>60 years) adults. These tasks uniquely combined bimanual control and altered dynamic conditions to assess age-related differences. The first task required bimanual coordination to control a virtual 2D arm over 400 center-out and return trials, targeting de novo motor learning. The second task involved unimanual reaching with the dominant hand, consisting of 200 trials in a null-field condition followed by 200 trials with object-like dynamic forces. The third task similarly began with 200 null-field trials and then introduced a viscous force field in the final 200 trials, with fast movements rewarded to encourage peak performance. This task also enabled comparison between dominant and non-dominant arms. All tasks detected age-related performance differences, with the viscous resistance task proving most sensitive to declines in movement speed, force generation, and response onset time. Scoring mechanisms that encouraged brisk performance amplified these effects. Across tasks, older adults generally moved more slowly, took longer to complete tasks, exerted lower peak forces, and had longer response onset times. However, some older participants performed comparably to younger individuals. In the third task, dominant arm performance consistently exceeded that of the non-dominant arm. These results demonstrate that robot-based tasks can sensitively quantify age-related sensorimotor decline and may offer valuable metrics for clinical assessment and monitoring.

**Significance Statement:** This study utilizes a robotic manipulandum, i.e., the vBOT, to assess the effect of aging. Through precisely controlled repeatable sensorimotor tasks, we quantify movement characteristics in both younger and older adults, providing clear insights into age-related motor decline. Our approach not only highlights the magnitude and variability of functional changes with age but also establishes rigorous and reliable quantitative benchmarks that could serve as critical tools for evaluating preventative therapies. In turn, this could guide the development of interventions aimed at preserving motor performance in older age.

**Highlights:** This study introduces a novel approach to assessing how aging affects movement dynamics using a robotic system. As populations age, understanding sensorimotor decline becomes increasingly crucial. Unlike conventional assessments, our method quantifies motor control metrics with precision, revealing previously undetectable, subtle age-related changes in movement efficiency, response onset times, and force generation. This framework establishes a rigorous benchmark for evaluating age-related decline, offering valuable clinical applications and informing the design of interventions to support older adults.

## 1. Introduction

Over the past century, life expectancy has shown a steady increase. In 1920, the global average at birth was just 36.4 years; by 2020, it had more than doubled to 72.5 years (Dattani et al., 2023). Due to this significant rise, combined with other factors, the population over the age of 60 is steadily expanding. By 2030, this age group is estimated to grow from 1 billion in 2019 to 1.4 billion, with projections suggesting it will further increase to 2.1 billion by 2050 (Grinin et al., 2023; WHO Global, 2024). This growth in the aging population introduces a range of challenges, particularly in the context of health and associated age-related changes in human motor control.

With increasing life expectancy and a growing population of individuals over 60 years of age, it would be helpful to develop quantifiable sensorimotor indicators that signal the onset of age-related decline in healthy older adults. Within-participant comparisons across age would be especially informative, as they avoid the confound of inter-individual variability in baseline ability. Identifying and measuring these early markers can equip health professionals with precise, objective, and reliable tools to assess sensorimotor function, decline and the extent of movement improvements resulting from therapy.

Pioneering work using robotic manipulanda, such as the MIT-Manus (Hogan et al., 1992) and Kinarm (Scott, 1999) devices, has significantly advanced the quantitative assessment of sensorimotor performance during upper-limb movements. These robotic systems precisely measure and control limb dynamics and kinematics, enabling detailed examination of motor control strategies, sensory integration, and adaptation. The MIT-Manus has been instrumental in characterizing impaired motor function post-stroke, revealing deficits in movement smoothness, accuracy, and force control, which correlate closely with clinical evaluations of motor impairment (Krebs et al., 2000; Krebs and Hogan, 2006). Similarly, the Kinarm robotic system has facilitated investigations into sensorimotor integration and proprioception, allowing researchers to isolate and quantify specific deficits in motor coordination and sensory processing after neurological injuries or disorders (Dukelow et al., 2010). Robotic manipulanda provide precise and objective measurements, overcoming limitations of subjectivity and limited sensitivity inherent in traditional assessments.

Robotic manipulanda are extensively utilized to assess motor deficits associated with aging and various neurological conditions, providing objective, sensitive metrics beyond traditional clinical assessments. The high precision and repeatability of robotic measurements (Scott and Dukelow, 2011) allow the early detection of subtle impairments in motor performance (Simmatis et al., 2017), such as characteristics of aging (Herter et al., 2014). Studies employing the Kinarm, for example, have identified age-related declines in proprioception, reaction time, and motor coordination, highlighting the subtle sensorimotor changes that can precede overt functional deficits in some older adults, although they do not necessarily progress to clinically meaningful impairment (Herter et al., 2014; Kitchen and Miall, 2019).

Additionally, robotic assessments have been helpful in evaluating the progression and severity of conditions like Parkinson’s disease and multiple sclerosis, facilitating early diagnosis, monitoring disease progression, and assessing therapeutic efficacy with greater accuracy and reliability (Bosecker et al., 2010; Coderre et al., 2010).

The research question we address is to develop and compare movement tasks using a robotic manipulandum that can indicate key aspects of movement performance, including the ability to cope with loads that exhibit viscous and inertial dynamics and the capacity to learn a novel bimanual coordination pattern. We aim to determine whether behavioral differences can be observed between younger and older participants across tasks that probe different aspects of dynamic control, reward, and learning, and to identify which tasks are most sensitive to revealing these age-related differences. In addition, a central aim of the study is to assess how younger and older adults adapt when exposed to newly introduced dynamic environments. By examining changes in movement metrics across both null-field and perturbed conditions, we explicitly evaluate age-related differences in motor adaptability.

To address these goals, the study uses three complementary experiments, each targeting a different facet of sensorimotor performance. Experiment 1 examines de novo bimanual coordination and learning of a novel kinematic mapping. Experiment 2 focuses on adaptation when an inertial load is introduced, assessing force production and deceleration control. Experiment 3 evaluates adaptability under a velocity-dependent viscous perturbation and the influence of performance incentives. Together, the three paradigms probe distinct but complementary dimensions of motor control. These experimental tasks used the vBOT robotic manipulandum (Howard et al., 2009), and we specifically compared performance between healthy young adults (under 35 years) and older adults (over 60 years).

## 2. Methods

To develop assessments suitable for future participants in a clinical setting, it was necessary to limit the duration of the tasks they would be required to perform. To ensure participants were not fatigued by the assessment procedure, we restricted the overall testing time to 30 minutes per assessment. Since the current study involves investigating the appropriateness of the assessments we were developing, we ran two experimental sessions per participant to collect as much data as possible. Because we included older participants in our study, we split the experiments into two groups so that each participant was required to complete only two experiments. This approach avoided fatiguing participants with long sessions, ensured that we did not ask for more than one hour of their time, and prevented the need for them to return for an additional session on a different day.

The two-session design consisted of either running two different experimental protocols on the same day with the right hand (Experiments 1 and 2) or running the same protocol first with the right and then with the left hand (Experiment 3). The only exception was a single participant who completed three experiments: first Experiment 3 with both hands, and then, two months later, Experiments 1 and 2. The two-month interval ensured that there was little, if any, practice effect between the viscous and mass paradigms that could influence performance. We note that in future clinical use, the goal would be to perform a single test session rather than multiple research-focused sessions.

Participants: We recruited a total of 32 right-handed participants, consisting of 16 young adults and 16 older adults. Participants were screened to exclude individuals with a history of neurological conditions, musculoskeletal disorders, or significant visual impairments that could affect performance. Half of the young adults and half of the older adults were randomly assigned to Experiments 1 and 2. Participants first completed Experiment 1, followed by Experiment 2. The remaining participants were assigned to Experiment 3. All but one of them completed it twice: once using the right arm and once using the left.

All participants were naïve to the objectives of the experiments and provided written informed consent prior to participation. Right-handedness was assessed using the Edinburgh Handedness Questionnaire (Oldfield, 1971), which confirmed that all participants were right-handed. All experiments followed the ethics protocol approved by the University of Plymouth’s Faculty Research Ethics and Integrity Committee.

### 2.1. Experiments

We developed three experiments that each investigated different aspects of human motor control:

Experiment 1: Learning to Control a 2D Kinematic Arm examined how participants learned to control a two-dimensional virtual arm using movements of both the left and right hands. The aim was to investigate how participants acquire novel forms of bimanual coordination and develop new kinematic mappings to perform center-out and out-to-center movements. This task also required the use of the participants’ ability to use cognitive strategies to master unfamiliar movement relationships, rather than relying on existing motor patterns.

Experiment 1 investigated 8 older adults (5 females and 3 males; mean ± SD age: 73.61± 6.24 years) and 8 younger adults (4 females and 4 males; mean ± SD age: 28.02 ± 2.92 years).

Experiment 2: Handle Mass Centre-Out Movement involved making unimanual, right-handed point-to-point movements, initially in an unloaded null-field and later with a simulated mass at the handle. The aim of this experiment is to investigate differences in behavior between younger and older adults when subjected to an inertial load, which simulates aspects of moving an object in the real world. This includes managing acceleration and then appropriately decelerating the inertial mass to avoid overshoot.

In Experiment 2, performance was investigated mainly reusing the same participants as in Experiment 1 (1 participant needed to be discarded due to problems in the mass task with stability and replaced with another). It involved 8 older adults (5 females and 3 males; mean ± SD age: 73.61 ± 6.24 years) and 8 younger adults (3 females and 5 males; mean ± SD : 28.69 ± 3.01 years).

Experiment 3: Handle Viscous Resistance Center-Out Movement, involved making unimanual point-to-point movements, initially in an unloaded null-field condition and later with simulated viscosity at the handle. The aim of this experiment is to again investigate movement involving significant force production, more akin to moving in water. It also includes providing a score for the number of times movements were made within a desired timeframe, thus offering a reward signal to motivate participants.

In Experiment 3, almost all participants ran the task twice, once with each hand. The cohort used to investigate the effect of aging on right-hand performance included 8 older adults (4 females and 4 males; mean ± SD age: 69.09 ± 2.97 years) and 7 younger adults (4 females and 3 males; mean ± SD age: 22.42 ± 2.61 years). Left-hand performance was examined in all 7 younger adults and a smaller subset of 7 older participants, since one older participant was not available (leaving 4 females and 3 males; mean ± SD age: 68.35 ± 2.27 years).

### 2.2. Apparatus

Fig. 1 illustrates the experimental setup used in this study to analyze upper limb reaching movements. Participants interacted with a pair of vBOT robotic manipulanda (Howard et al., 2009). Each was equipped with a robotic handle that incorporated an activation switch, which participants needed to press during a trial to activate the controller and allow the experiment to proceed. During the experiment, participants were seated in a comfortable, cushioned chair and secured with two shoulder straps to reduce body movement. For the unimanual experiments, a participant’s relevant forearm rested on an air sled, ensuring low-friction movement constrained to a 2D plane. For the bimanual experiments, both forearms were supported by air sleds. Throughout all experiments, participants could not directly observe their arms or hands; instead, veridical visual feedback was provided using a 2D virtual reality system, which displayed visual elements such as circles representing hand positions, target locations, and start points within the workspace.

**Figure 1.**
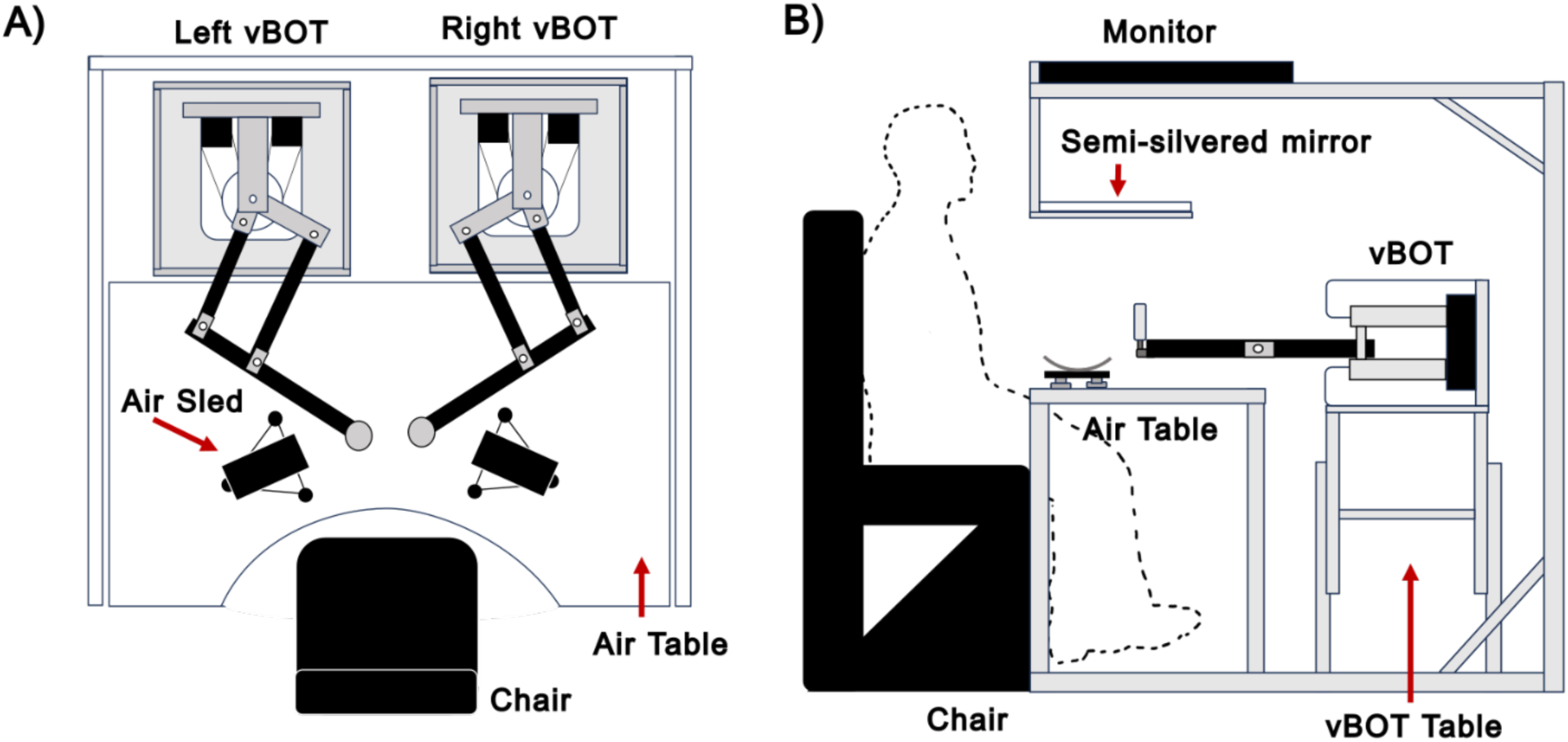
Schematic Illustration of the vBOT Robotic Manipulandum. **A:** Plan view of the vBOT robotic manipulandum setup. **B:** Side view of the vBOT robotic manipulandum setup.

Handle positions were measured using optical motor encoders, sampled at 1kHz, while torque-controlled motors generated endpoint forces. A force transducer (Nano 25; ATI), mounted beneath the left vBOT handle, measured the forces applied by participants during unimanual trials. These force signals were low-pass filtered at 500 Hz using 4th-order analog Bessel filters prior to digitization at a 1kHz sampling frequency. Force and position data were stored for offline MATLAB analysis.

### 2.3. Experiment 1: Learning to Control a 2D Kinematic Arm

Participants used both arms (left and right) to control a cursor in a center-out and out-center task by holding the handles of two robotic manipulanda. The cursor position corresponded to the endpoint of a 2D virtual planar arm, and the participants’ forward and backward handle movements controlled its joint angles, This is a modification of a paradigm used in a study to investigate *de novo* learning (Howard and Alvarez-Hidalgo, 2025). The experimental setup and design are shown in Fig. 2.

**Figure 2.**
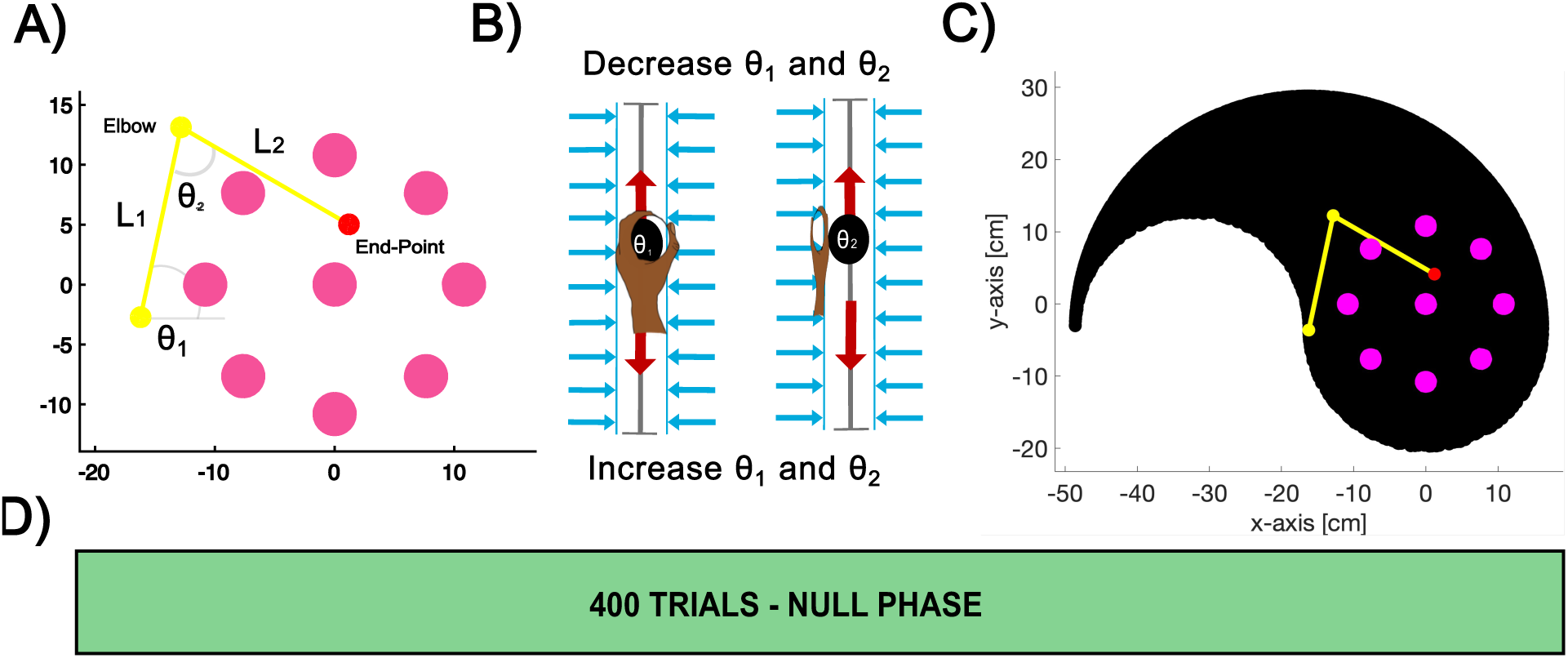
Experiment 1: Learning to Control a 2D Kinematic Arm. **A:** Experimental setup. Eight pink target points were arranged in a circular pattern at specific angular positions. The pink center point represents the starting position during center-out trials, and the target during out-to-center trials. A red cursor indicates the endpoint of a virtual 2D robotic arm with its links displayed in yellow. **B:** Dual-arm control mechanism. The left hand controls the joint angle θ_1_, and the right hand controls the joint angle θ_2_. **C:** The robotic arm’s reachable workspace is shown in black, illustrating the reachability of the virtual arm endpoint when driven by the dual-arm control positions. **D:** Trial design. Each participant completed a total of 400 trials during the null-field phase.

#### Virtual 2D Arm

Participants interacted with a simulated two-link robotic arm displayed in yellow, whose endpoint was represented by a red cursor. As shown in Fig. 2A, eight possible pink target points (radius: 1.0 cm) were arranged in a circular pattern at a radius of 10.8 cm from a central pink point (radius: 1.0 cm) and located at workspace origin (0,0). Targets were positioned at 0°, 45°, 90°, 135°, 180°, 225°, 270°, and 315°. The center point also served as the starting position for center-out trials and as the target for out-center trials.

Participants controlled the virtual 2D arm by grasping two robotic handles, each linked to a specific joint angle of the virtual 2D arm. The left handle controlled joint angle *θ₁*, and the right handle controlled joint angle *θ₂*, as depicted in Fig. 2B. Vertical hand movements adjusted these joint angles: upward movements decreased the angles, while downward movements increased them. To constrain hand movements to the y-axis, participants operated within two vertical channels (Scheidt et al., 2000).

#### Trials Procedure

The experiment involved two types of movements: center-out movements and out-to-center movements. In center-out movement trials, participants started at the central position and moved the red cursor to one of the eight peripheral targets. In contrast, in out-to-center movement trials, participants started at a peripheral target and moved the red cursor back to the central position. Each trial began with the participant pressing both activation switches on the back of the robotic handles. We did not implement a familiarization period in which trial data were excluded, because we wished to minimize the overall length of the experimental session and because initial behavior is itself indicative of a participant’s movement strategy and performance.

At the start of each center-out trial, the potential peripheral targets were first displayed in pink, while the central position appeared in grey. Participants then positioned the arm’s endpoint cursor at the starting location. Once the cursor remained stationary at this location for 500 ms, the vBOT emitted an auditory cue (a beep). Simultaneously, one of the eight target locations was selected pseudorandomly and turned yellow, signaling the direction in which the participant should perform the trial. The virtual arm also appeared at this point. Participants were instructed to move the cursor into the target as quickly as possible and to remain stationary once this had been achieved. As the cursor moved out of the start location, the central grey point disappeared. The trial ended when the red cursor entered the target and stopped moving (speed < 0.01 cm s⁻¹), at which point the virtual arm also disappeared. They were not scored on their performance, and no reward was provided.

After completion of a center-out movement trial, participants performed an out-to-center trial, which returned the cursor to the central position. The procedure was similar to the center-out trials, except the movement began at a peripheral location, and the central position turned yellow to indicate it as the target.

#### Experimental Block Design

As illustrated in Fig. 2D, the experiment consisted of 400 trials, divided into 25 blocks of 16 trials each. Each block contained eight center-out trials and eight out-to-center trials. The peripheral target was assigned pseudo-randomly within each block. No formal breaks were scheduled, but participants were informed that they could pause the experiment at any time by releasing the handle switch; the experiment resumed when they closed it again.

#### Mechanical Channels

Participants’ hand movements were constrained to two distinct vertical channels, each corresponding to one hand. These channels were aligned along the y-axis with a length of 30 cm (chanLen). Both channels were offset along the x-axis by ±10 cm from the center of the workspace, with the midpoints of the sliders located at (−10, 0) for the right hand and (+10, 0) for the left hand.

These linear channels were characterized by a stiffness value of 4,000 N.m⁻¹ (*k_chan_*) and a damping coefficient of 30 N·s·m⁻¹ (*b_chan_*). Forces experienced by each hand within the channels were determined by the following equation, which relates the force to the deviation of the handle in the x-axis from the channel position (xₕ - x₀, where xₕ is the handle’s x-position and x₀ is the x-location of the channel) and to the x-axis velocity:

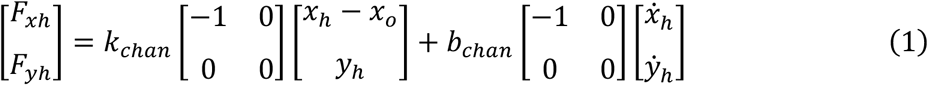

#### Control of the Simulated Arm

The vertical position (*y*) of each handle was linearly scaled to map to the corresponding joint angle (𝜃). This mapping was defined as:

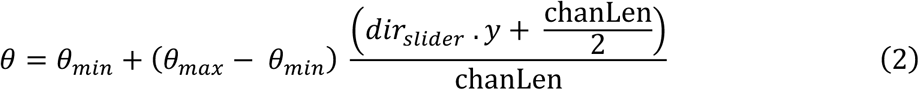

where chanLen is the length of the channel, and 𝑑𝑖𝑟*_silder_* sets the direction in which the slider acts (and has a value of 1 or -1).

For the left handle, the vertical position (*y_h1_*) was mapped to a joint angle (𝜃*_1_*), within the range of 𝜃*_min_* = 0 radians and 𝜃*_max_* = π radians. Similarly, the vertical position (*y_h2_*) of the right handle was mapped to a joint angle (𝜃_2_), within the range of 𝜃*_min_* − π radians and 𝜃*_max_* = 0 radians.

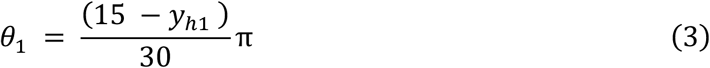

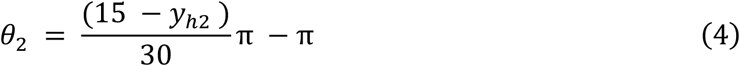

This allowed the y-position of the handles to be converted into joint angles within their respective ranges, resulting in the reachable workspace shown in black on Fig. 2C.

#### Simulating Forward Kinematics

To compute the endpoint and visualize the configuration of the 2D arm, its forward kinematics were used. The simulated arm link lengths *l_1_* and *l_2_* were both 16.2 cm long. By employing trigonometric analysis, the positions of the elbow and end effector in Cartesian coordinates can be described using matrix equations:

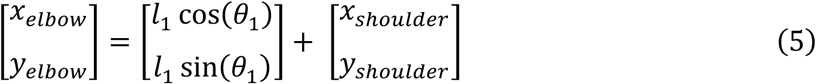

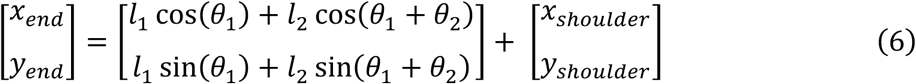

The shoulder position (-16.2, -3.6) was located slightly below the central start location and to its left just outside the circle on which the targets are located.

### 2.4. Experiment 2: Handle Mass Center-Out Movement

In this experiment, participants performed center-out movements to one of eight targets. The experiment assessed their performance under two conditions: a null field allowing free movement, and a condition simulating the effect of moving a 5 kg mass located at the handle. The latter tested their ability to generate movements requiring substantial force production. The experimental design is summarized in Fig. 3.

**Figure 3.**
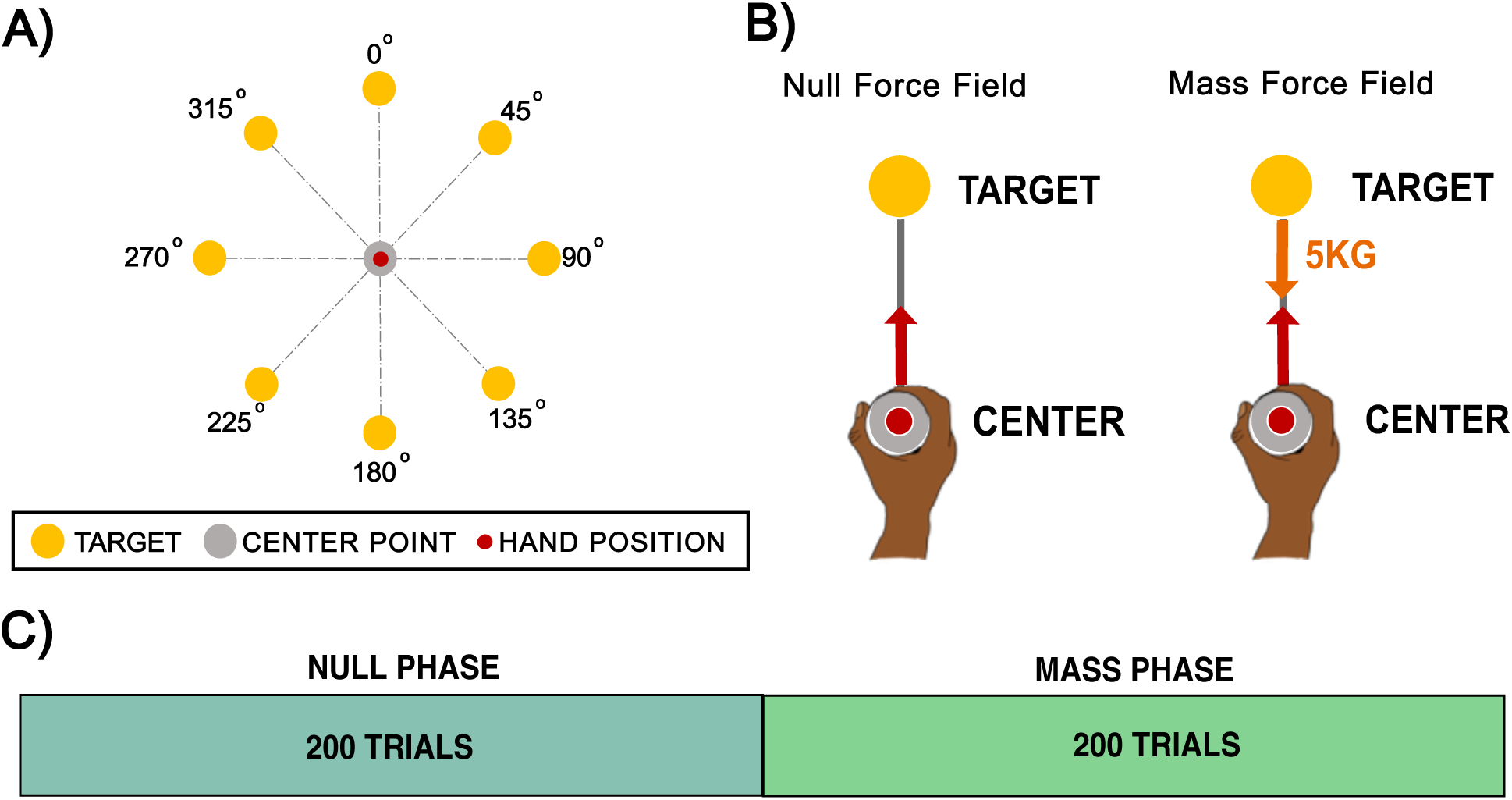
Experiment 2: Handle Mass Center-Out Movement. **A:** Experimental setup. Eight yellow targets were arranged at specific angular locations. A grey center point represented the starting position, while a red cursor indicated the participant’s hand location. **B:** Experimental phases. Participants experienced either a null-field, allowing free movement within the workspace, or the effect of a mass. The latter introduced inertial resistance by simulating the sensation of moving a 5 kg point mass around the workspace. **C:** Experimental conditions. The experiment consisted of 400 trials in total: 200 trials in the null-field phase, followed by 200 trials in the point mass phase.

#### Experimental Setup

Participants were instructed to perform center-out movements toward one of eight possible yellow targets positioned in a circular pattern around a grey center point. As depicted in Fig. 3A, the starting position was represented by a central grey circle (radius: 1.25 cm) at the center of the workspace (0,0), and a red cursor (radius: 0.5 cm) displayed the hand’s position in the workspace. Additionally, eight yellow target circles (radius: 0.5 cm) were arranged in a circular pattern at a radius of 12 cm from the center, located at angular positions of 0°, 45°, 90°, 135°, 180°, 225°, 270°, and 315°. Only one target appeared on the screen at a time.

Participants controlled the red cursor by holding the left vBOT robotic handle with their right hand. At the start of each trial, participants viewed an empty screen and grasped the left vBOT robotic handle. The vBOT guided their hand to the designated center position. After remaining stationary at this location for 300 ms, an auditory beep signaled the beginning of the trial. At the same time, the hand position cursor, as well as a single yellow target, appeared on the screen. At this point, participants were required to move their hand from the center position toward the displayed target. The trial concluded when the participant successfully reached the target.

#### Experimental Block Design

Movements were undertaken under two conditions. During this center-out movement, participants encountered either a null-field or a 5 kg mass (see Fig. 3B). Each experiment consisted of 400 trials, with the first 200 conducted in a null-field phase and the remaining 200 in an endpoint mass phase. Approximately midway through the experimental condition, participants were given a 10-second rest. However, participants could take a break at any time simply by releasing the activation switch. The timeline of trials is shown in Fig. 3C.

#### Simulating Mass

To ensure a stable simulation, moving the mass was modeled as being connected to the hand cursor via a stiff spring. Additionally, viscous damping was incorporated in the simulation, to enhance stability. At the start of the simulation, the mass’ position is initialized to match the current location of the hand cursor. The mass’ velocity, acceleration, and the forces acting on it are all set to zero.

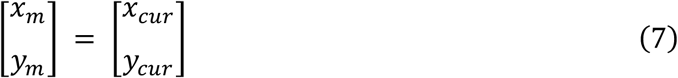

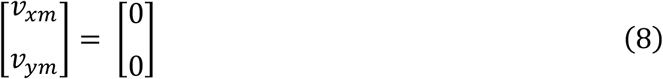

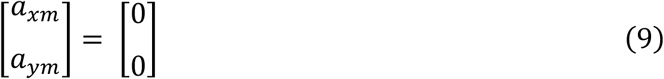

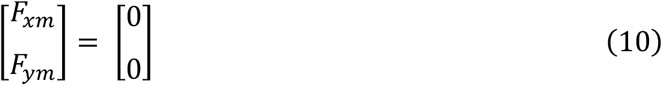

Within the main simulation update loop, which operates at a nominal frequency of 1 kHz, the motion of the mass is updated in real time. At each time step, the position of the mass is updated using Euler integration based on its velocity:

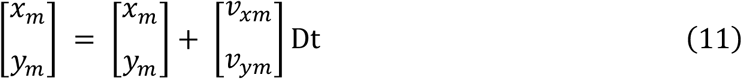

Here, Dt represents the timestep since the last update, typically 1 ms. The mass’ acceleration is then calculated using the following equation:

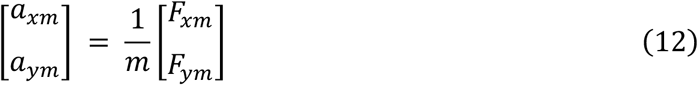

where m = 5 kg is the mass and 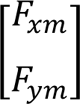 is the force acting on it. Using Euler integration, the velocity of the mass is updated based on its acceleration:

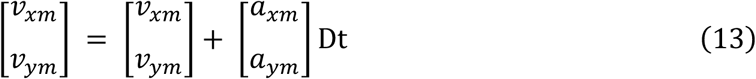

The force acting on the mass is determined by the extension of the spring, which connects the mass to the endpoint of the arm. The spring has a stiffness constant of *k_mass_* = 2000 N.m^-1^:

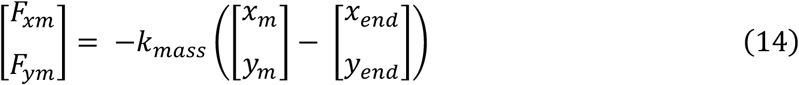

The endpoint velocity, which is directly available from the handle velocity, is used to calculate an additional viscous damping force for further stability enhancement. The viscous damping force is given by:

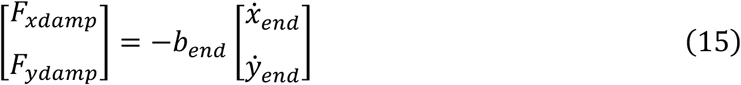

Here, the damping viscosity coefficient (*b_end_*) is 10 N.s.m^-1^. Finally, the spring force (Eq. 14) and damping force (Eq. 15) are combined to compute the total force experienced at the handle.

### 2.5. Experiment 3: Handle Viscous Resistance Center-Out Movement

This experiment was similar to Experiment 2. Participants again performed center-out movements to one of eight targets. However, after the initial 200 trials, during the second phase of the experiment, instead of introducing a 5 kg simulated point mass at the handle, a viscous force field that resisted hand movement was introduced. This resistance was proportional to the velocity of movement, producing a sensation similar to moving underwater. The experimental design is illustrated in Fig. 4.

Experiment 3 was first carried out with their right hand. To compare dominant and non-dominant arm movement performance, some participants also repeated the experiment with their left hands.

#### Viscous Force Field

To implement viscous resistance in the direction of movement, we used the force equation:

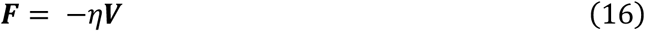

Where ***F*** and ***V*** are vector quantities, and η is the viscous-field constant with a value of 0.3 N·s·m⁻¹.

**Figure 4.**
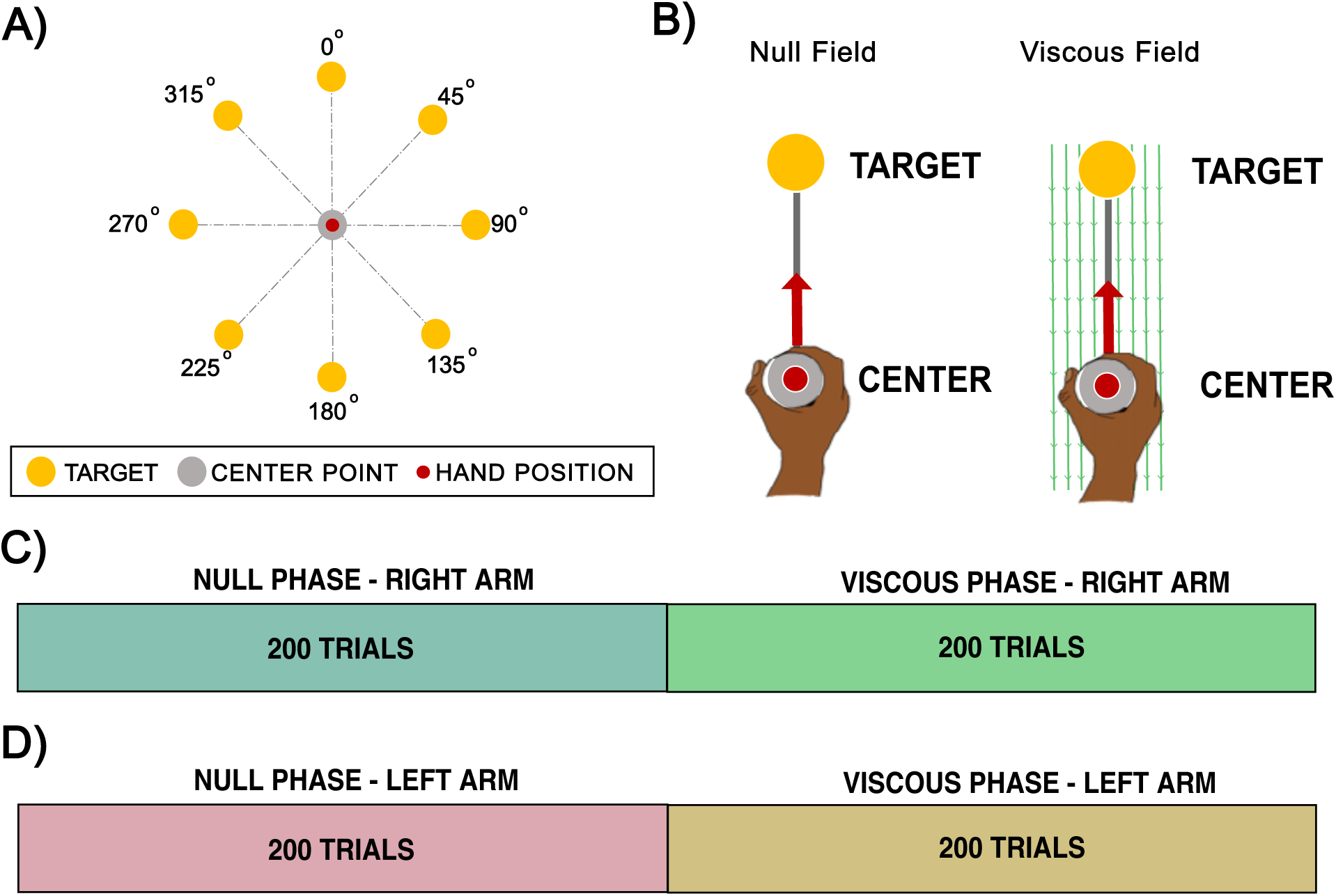
Experiment 3: Handle Viscous Resistance Center-Out Movement. **A:** Experimental setup. Eight yellow targets were arranged at specific angular locations. A grey center point represented the starting position, while a red cursor displayed the participant’s hand location. **B:** Experimental phases. Participants experienced either a null-field, allowing free movement within the workspace, or a viscous-field, which introduced resistance. **C:** Right-Hand Experimental Condition. Participants performed the experiment with their right arm for all 400 trials: 200 trials in a null-field phase, followed by 200 trials in a viscous-field phase. **D:** Left-Hand Experimental Condition. Participants repeated the same experimental procedure performed with their right hand, but this time using their left hand.

#### Experimental Block Design

Each condition consisted of 400 trials, with the first 200 conducted in a null-field phase and the remaining 200 in a viscous force-field phase. Approximately midway through the experimental condition, participants were given a 10-second rest period. Participants could also take a break at any time simply by releasing the activation switch. The timeline of trials to assess right-hand performance is shown in Fig. 4C. In this experiment, participants received a score reflecting the number of times they achieved a brisk movement (duration < 600 ms), thereby encouraging them to move faster than they otherwise might have done. A. Movements shorter than 450 ms were classified as too fast and were not rewarded.

#### Dominant versus Non-Dominant Arm Usage

To assess how dominant versus non-dominant arm use affects performance on movement characteristics and performance, each participant performed the experimental paradigm twice, with a short break between experimental sessions: once using their right arm and once using their left arm (refer to Fig. 4D). This also provided an effective means to examine the robustness of the movement analyses, as it permitted intra-participant comparisons that were unaffected by the different levels of performance that occur across participants.

## 3. Data Analysis

The experimental data was processed offline using MATLAB R2024a. For the de novo arm experiment (Experiment 1), we examined the trajectory of the cursor, which corresponded to the endpoint of the virtual arm. For the mass experiment (Experiment 2) and the viscous experiment (Experiment 3), we also examined the trajectory of the cursor, which corresponded to the location of the robot handle.

Each trial began with an auditory cue (a beep). Movement onset here was defined as the moment the hand exceeded a distance of 1.25 cm from the starting position, and its speed surpassed 5 cm·s⁻¹. Movement termination was defined as the moment the hand entered within 1.25 cm of the target and its speed dropped below 5 cm·s⁻¹.

### 3.1. Hand Trajectory Plots

For the de novo arm experiment (Experiment 1) we plot the mean ± standard error (SE) of movement trajectories over the first and final blocks of trials to provide an intuitive visualization of participants’ movement behavior.

### 3.2. Performance Metrics

We quantified kinematic accuracy with six metrics:

#### Absolute Maximum Perpendicular Error (AMPE)

For each trial, we computed the absolute maximum perpendicular deviation of the movement trajectory from the ideal straight line connecting the start and target cursor locations in extrinsic space. By taking the absolute value, this metric avoids positive and negative deviations from canceling each other out across trials and provides a clear overall measure of how accurately participants followed a straight path from the starting point to the target.

#### Movement Duration

The time taken to complete a movement was calculated as the difference between the movement onset and termination times.

#### Movement Length

The total path length was determined by summing the distances between consecutive (x, y) cursor positions along the trajectory, from movement start to movement end.

#### Response Onset Time

This is the time elapsed between the cue to start moving and the moment the cursor exits the home position with a speed exceeding 5 cm·s⁻¹. This measure reflects both the initiation of movement (reaction time) and the initial acceleration required for the cursor to reach the detection threshold.

#### Maximum Speed

The highest speed attained during the movement.

#### Maximum Force

The peak force exerted in the direction of movement.

We note that both Absolute Maximum Perpendicular Error (AMPE) and Movement Length measures quantify aspects of how a trajectory deviates from a straight line, but they capture different properties of that deviation. AMPE reflects the single largest lateral deviation from the ideal straight path, whereas Movement Length reflects the accumulated deviation across the entire trajectory. As a result, the two metrics are not equivalent. For example, a path with substantial jitter will produce an increased Movement Length but may still have a relatively small AMPE because no individual deviation is large. Conversely, a single pronounced deviation would increase AMPE without greatly affecting the total path length. These differences show that each metric provides distinct and complementary information about movement quality.

### 3.3. Statistical Analysis

Statistical analyses were performed using JASP software (JASP Team, 2024). When appropriate, Mauchly’s test was applied to assess the assumption of sphericity. If this assumption was violated, Greenhouse-Geisser corrections were used to adjust the degrees of freedom. The significance level (*α*) was set at 0.05 for all tests. Omega squared (*ω²*) was reported as a measure of effect size, indicating the proportion of variance in the dependent variable explained by the independent variables.

Critical Phases and Statistical Comparison: For each experimental condition, we identified three critical phases of the trial sequence:

1. The initial batch of trials, representing early performance.
2. The final batch of trials, representing stabilized performance.
3. The condition-change phase, corresponding to transitions where the dynamic properties of the task were altered and switched from a null field phase to exposure phase, for example, when movements were made without any additional dynamics applied to the handle, or during exposure phases when additional dynamics were introduced, such as simulated mass (Experiment 2) or viscosity (Experiment 3).

These phases were selected because they capture points at which behavior is expected to differ most strongly: initial exposure, stable performance, and adaptation to new task dynamics. For each metric, repeated-measures ANOVAs were conducted to compare performance across these phases within each experimental condition. We also performed statistical tests across age groups to examine differences due to ageing.

When a significant main effect was observed, where appropriate we proceeded with post hoc comparisons using the Holm-Bonferroni correction. Bonferroni-corrected p-values are indicated as *p_Bonf_*.

Note that block size differed by experiment; references to ‘exposure blocks’ correspond to eight-trial blocks unless otherwise specified. To compare metrics across the conditions, of age (younger versus older participants and the use of the right versus left hand), separate ANOVAs were conducted on the initial exposure metrics (defined as the mean of the first 25 exposure blocks) and the final exposure metrics (mean of the last 25 exposure blocks).

## 4. Results

### 4.1. Experiment 1: Learning to Control a 2D Kinematic Arm

The first experiment was a novel bimanual kinematic arm task in which participants controlled the endpoint of a simulated two-dimensional (2D) arm to complete a center-out task. Their left- and right-hand positions, restricted to vertical movements, determined the arm’s joint angles. The experiment consisted of a total of 400 trials (200 center-out and 200 out-center).

Fig. 5 shows the mean and standard error (SE) of both outwards and inwards endpoint movement trajectories of the virtual arm at two critical stages, for both younger and older participants. Panel A illustrates the first 16 trials of the younger participants, where movements are highly erratic, reflecting exploratory behavior as they attempt to gain control of the arm’s endpoint. By the end of the experiment, as shown in Panel B, trajectories become much straighter, and deviations from the mean direction are markedly reduced. Similar improvements are observed in older participants. Panel C displays the first 16 trials for the older group, also showing highly erratic movements. However, by the end of the 400 trials, their performance improves significantly as well. Qualitatively, younger participants appear to perform slightly better than older participants, in terms of the straightness of their movements. We also note that for both sets of participants, inward movements were straighter than outward movements, no doubt arising from the fact that it was easier to move back to the central location than outwards to one of many peripheral targets.

We hypothesize that inward movements were easier to generate than outward movements for two main reasons. First, outward movements required reaching to one of eight pseudo-randomly selected peripheral targets. In contrast, inward movements were always made towards a single central location, which was visited eight times more frequently than any peripheral target. As a result, the centre location provided greater opportunity for learning, consolidation, and recall due to its high repetition rate. Second, the inward target location could be supported by short-term sensorimotor memory, as each centre-directed movement was consistently preceded by an outward movement and followed by a return to the centre, forming a highly predictable out–in movement sequence.

**Figure 5.**
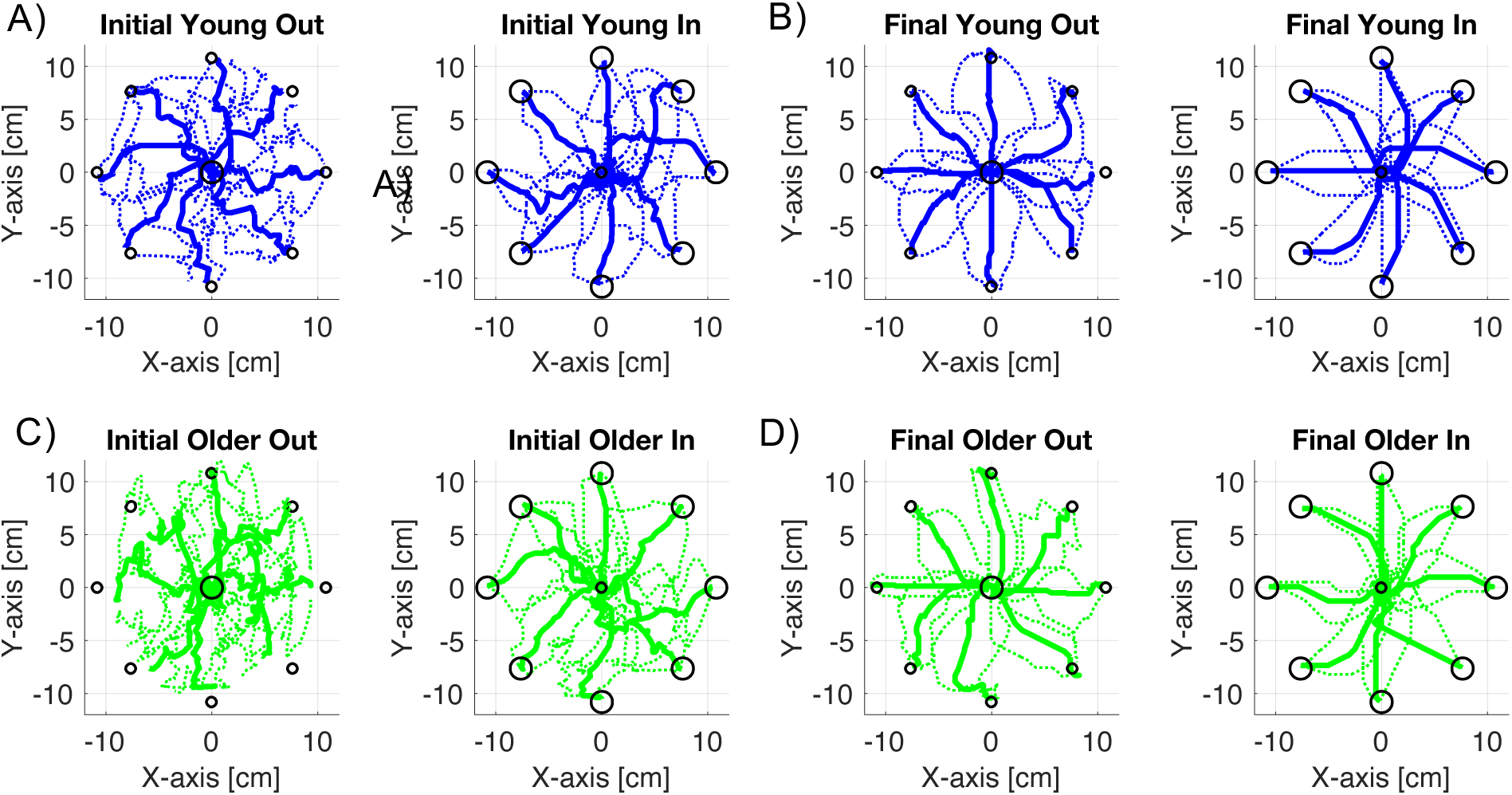
Plots showing the mean ± standard error (SE) of virtual arm endpoint trajectories over 16 trials for outward and inward movements: A: Under-35 years, initial trials (1 - 16); B: Under-35 years, final trials (385–400); C: Over-60 years, initial trials (1–16); D: Over-60 years, final trials (385–400).

Behavioral trends related to age were evident in the kinematic arm task data. Younger adults, in general, took less time to complete each trial and made straighter movements in extrinsic space. These quantitative indicators of motor performance across age groups may serve as a means for evaluating the effectiveness of future rehabilitation strategies.

Fig. 6 displays movement metrics used to quantify the performance of the endpoint movement of the virtual arm. Each metric reflects the mean of eight consecutive trials (corresponding to four center-out trials and their four associated out-to-center movements) and is plotted as a function of blocks of eight trials. Across all metrics, there is a general tendency for performance to improve with practice. Panel A shows that movement duration starts off high for both younger and older participants but decreases considerably by the end of the experiment. Younger participants exhibit consistently shorter movement durations than older participants. Panel B presents the absolute maximum perpendicular error (AMPE) across blocks, with a general downward trend. Younger participants tend to show slightly lower AMPE values compared to older participants. Panel C depicts response onset time - defined as the interval between the cued start of a trial and measured movement initiation. Younger participants consistently demonstrate shorter response onset times. Panel D shows the maximum movement speed as a function of blocks, revealing a slight overall increase with practice. Younger participants move faster than older participants on average. Panel E displays trajectory path length, which is consistently shorter in younger participants across all trial blocks.

#### Comparison of Results within Age Groups

Repeated-measures ANOVAs were used to examine whether there were any significant variations in the measured metrics across the initial and final two blocks (32 trials) for both younger and older participants. They revealed some significant improvements in task performance across training for both younger and older participants. Detailed results are summarized in Table 1.

Duration, Absolute Maximum Perpendicular Error (AMPE), Response Onset Time, and Path Length all significantly decreased from the initial to the final blocks of training, indicating enhanced movement efficiency and response preparation over time. In the younger group, all four metrics showed marked improvements with large effect sizes, such as ω² = 0.643 for Duration. Similarly, the older group demonstrated significant gains, with particularly strong effects observed for Response Onset Time (F = 107.303, p < 0.001, ω² = 0.664). Overall, the results demonstrate clear learning effects within both age groups, with improvements across multiple performance metrics except for Peak Speed.

**Table 1.** Repeated-measures ANOVA results for each age group across the entire experiment: Separate repeated-measures ANOVAs were conducted for each performance metric to assess overall changes across the full duration of the task within each age group. Reported values include F-statistics, p-values, and effect sizes (ω²). Significant p-values are shown in bold.

| AGE GROUP | METRIC | F | P | $\omega^2$ |
| --- | --- | --- | --- | --- |
| Younger | Duration | 35.808 | **< 0.001** | 0.643 |
|  | AMPE | 20.611 | **0.003** | 0.460 |
|  | Response Onset Time | 22.820 | **0.002** | 0.530 |
|  | Peak Speed | 0.014 | 0.909 | 0.000 |
|  | Path Length | 25.384 | **0.001** | 0.546 |
| Older | Duration | 44.317 | **< 0.001** | 0.741 |
|  | AMPE | 39.080 | **< 0.001** | 0.546 |
|  | Response Onset Time | 107.030 | **< 0.001** | 0.664 |
|  | Peak Speed | 0.121 | 0.738 | 0.000 |
|  | Path Length | 19.747 | **0.003** | 0.482 |

#### Comparison of Results Across Age Groups

To investigate age-related differences in performance, we compared average values for each metric between younger and older participants using an ANOVA, analysing the first half of trials (1–200) and the second half (201–400) separately. Detailed results are summarized in Table 2.

During Phase 1, no significant age-related differences were observed, though Duration showed a trend toward significance with a moderate effect size (ω² = 0.131). In Phase 2, significant differences were found in Duration (p = 0.008, ω² = 0.343) and Response Onset Time (p = 0.041, ω² = 0.202), indicating that older participants performed slower and responded more slowly in the later stage of the task. No differences were found in AMPE, Peak Speed, or Path Length.

**Table 2.** ANOVA results comparing younger and older participants within each experimental phase: Separate one-way ANOVAs were conducted for each performance metric during Phase 1 (P1, trials 1–200) and Phase 2 (P2, trials 201–400). Reported values include F-statistics, p-values, and effect sizes (ω²). Significant p-values are shown in bold.

| PHASE OF EXPERIMENT | METRIC | F | P | $\omega^2$ |
| --- | --- | --- | --- | --- |
| P1 | Duration | 3.419 | 0.086 | 0.131 |
|  | AMPE | 0.186 | 0.673 | 0.000 |
|  | Response Onset Time | 1.838 | 0.197 | 0.050 |
|  | Peak Speed | 0.679 | 0.424 | 0.000 |
|  | Path Length | 0.997 | 0.335 | 0.000 |
| P2 | Duration | 9.367 | **0.008** | 0.343 |
|  | AMPE | 0.296 | 0.595 | 0.000 |
|  | Response Onset Time | 5.038 | **0.041** | 0.202 |
|  | Peak Speed | 0.565 | 0.465 | 0.000 |
|  | Path Length | 0.980 | 0.339 | 0.000 |

**Figure 6.**
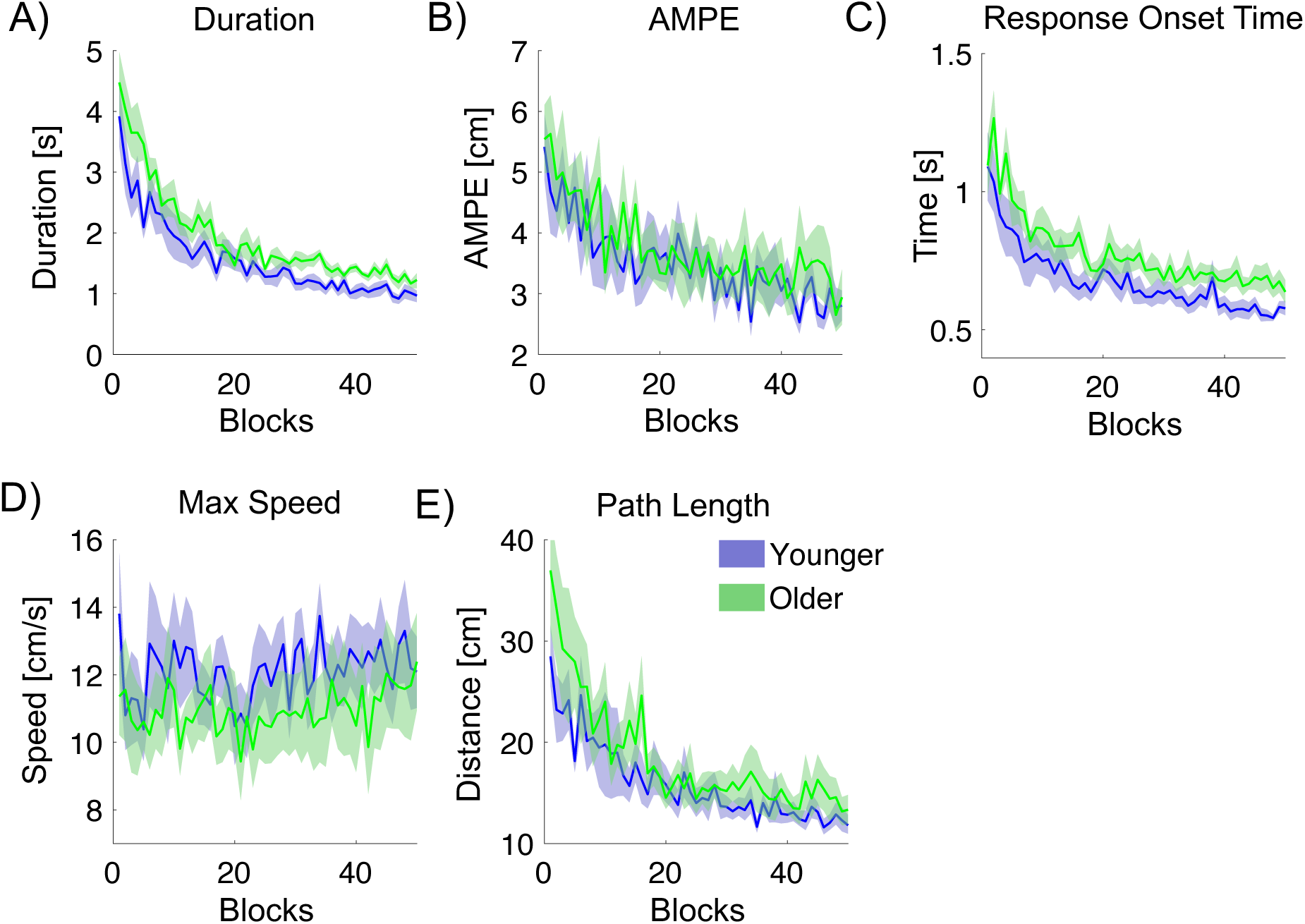
Movement metrics quantifying the trajectory at the endpoint of the virtual arm, calculated across center-out and out-to-center trials, and plotted against blocks of 8 trials. Younger participants are plotted in blue, and older participants are plotted in green. **A:** Duration of movement. **B:** Absolute maximum perpendicular error (AMPE). **C:** Response Onset Time. **D:** Maximum trajectory speed. **E:** Trajectory path length.

### 4.2. Experiment 2: Handle Mass Center-Out Movement

The second experiment was a right-hand center-out task involving two field conditions: a null field and a simulated endpoint mass. Participants made 12 cm movements from a central starting position to one of eight targets in each of two 200-trial phases.

The conditions were designed to investigate performance during unloaded and loaded movements. Behavioral trends related to age were evident: younger adults typically made faster, shorter-duration movements, generated greater peak forces in the direction of movement, and exhibited quicker response onset times.

Fig. 7 shows the performance of younger and older participants across several metrics quantifying hand trajectories during the mass center-out task. Compared to the older participants, younger participants take less time to complete the movement, have a faster response onset time, move with a higher peak speed, generate more force in the direction of movement, but have longer path lengths. However, AMPE is quite similar across the groups.

#### Comparison of Results within Age Groups

Repeated-measures ANOVAs were used to examine the significance of differences in the measured metrics between pairs of blocks of 8 trials (16 trials in total) located at the initial null exposure, final null exposure, initial mass dynamic exposure, and final mass dynamic exposure. Post hoc comparisons were conducted to further assess significant differences between these phases. Detailed results are summarized in Table 3.

Significant phase effects were observed in several performance metrics for both age groups. Bonferroni-corrected post hoc tests showed that the introduction of the mass dynamic (P_Bonf_ 2) led to consistent performance changes in AMPE, Peak Speed, Peak Y-Force, and Path Length for younger participants, and in Duration, AMPE, Response Onset Time, and Path Length for older participants. Adaptation across the mass dynamic phase (p_Bonf_ 3) was evident primarily in AMPE and Path Length in both groups. Only older participants exhibited significant changes in Response Onset Time. Path Length showed the largest effect sizes in both groups, indicating strong sensitivity to dynamic perturbation.

For younger participants, AMPE decreased significantly by the end of the null blocks, increased with the introduction of the mass dynamic, and decreased again by its end. Duration and Peak Speed increased significantly with the mass dynamic but did not change significantly otherwise. Peak Y-Force and Path Length also increased significantly with the mass dynamic; Path Length then decreased significantly by the end of this phase. Response Onset Time showed no significant changes.

For older participants, Duration decreased significantly during the null field phase, increased with the mass dynamic, and decreased again by its end. AMPE and Response Onset Time both increased significantly with the mass dynamic and decreased significantly by its end. Peak Y-Force and Path Length increased significantly with the mass dynamic, and Path Length also decreased significantly by the end of this phase. Peak Speed showed no significant changes.

Compared to younger participants, older participants showed significant changes in Response Onset Time and Duration across all phases, indicating a broader sensitivity to the mass dynamic. In contrast, younger participants showed no significant changes in Response Onset Time and more selective effects, with AMPE showing the clearest pattern of adaptation.

There is evidence that younger adults can typically generate higher maximal muscle forces and exhibit faster force development. However, the unloaded point-to-point reaching tasks used here require very little force to implement. It is therefore more likely that the observed differences arise from slower planning, execution, and feedback processes in older adults, rather than solely from reduced force-generation capacity.

**Table 3.** Repeated-measures ANOVA results for each age group across task phases: Each metric was analyzed using repeated-measures ANOVAs to assess changes across three task transitions: (1) from the initial to final null blocks, (2) from the final null blocks to the introduction of the mass dynamic, and (3) from the start to the end of the mass dynamic blocks. The table presents F-statistics, p-values, effect sizes (ω²), and Bonferroni-corrected post hoc p-values for each comparison. Significant p-values are shown in bold.

| AGE GROUP | METRIC | F | P | $\omega^2$ | P<sub>BONF</sub> (1) | P<sub>BONF</sub> (2) | P<sub>BONF</sub> (3) |
| --- | --- | --- | --- | --- | --- | --- | --- |
| Younger | Duration | 21.168 | < **0.001** | 0.539 | 0.244 | < **0.001** | 0.077 |
|  | AMPE | 18.210 | < **0.001** | 0.493 | 0.009 | **0.017** | **0.003** |
|  | Response Onset Time | 5.227 | **0.007** | 0.165 | 0.158 | 0.118 | 0.914 |
|  | Peak Speed | 6.705 | **0.002** | 0.161 | 1.000 | < **0.001** | 0.094 |
|  | Peak Y-Force | 26.530 | < **0.001** | 0.608 | 1.000 | **0.011** | 1.000 |
|  | Path Length | 40.052 | < **0.001** | 0.649 | 0.305 | < **0.001** | **0.001** |
| Older | Duration | 17.697 | **< 0.001** | 0.398 | **0.030** | **0.001** | **0.006** |
|  | AMPE | 9.258 | **< 0.001** | 0.384 | 0.155 | **0.009** | **0.017** |
|  | Response Onset Time | 6.113 | **0.004** | 0.129 | 0.947 | **0.026** | **0.003** |
|  | Peak Speed | 5.860 | **0.005** | 0.058 | 0.087 | 0.124 | 1.000 |
|  | Peak Y-Force | 20.825 | **< 0.001** | 0.519 | 1.000 | **0.013** | 0.663 |
|  | Path Length | 55.636 | **< 0.001** | 0.776 | 0.310 | **< 0.001** | **0.001** |

#### Across Age Groups Statistical Analyses

To investigate age-related differences in performance, we compared average values for each metric between younger and older participants using an ANOVA, analysing the first half of trials (1–200) and the second half (201–400) separately.

In Phase 1, significant age-related differences were observed in Duration, Peak Speed, and Path Length (all p < 0.05), with moderate to large effect sizes. Differences in Response Onset Time and Peak Y-Force approached significance. In Phase 2, the Response Onset Time did not reach significance (p = 0.050), but showed a moderate effect size (ω² = 0.184). Detailed results are summarized in Table 4.

In the null field phase movement duration is significantly shorter in the younger participants (*p* = 0.031), who also exhibit higher peak speed (p=0.032) and longer path lengths (*p* = 0.014). No other metrics showed a statistically significant difference between the two age groups.

**Table 4.** ANOVA results comparing younger and older participants within each experimental phase: Separate one-way ANOVAs were conducted for each metric within Phase 1 (P1, trials 1–200) and Phase 2 (P2, trials 201–400). Reported values include F-statistics, p-values, and effect sizes (ω²). Significant p-values are shown in bold.

| PHASE OF EXPERIMENT | METRIC | F | P | $\omega^2$ |
| --- | --- | --- | --- | --- |
| P1 | Duration | 5.779 | **0.031** | 0.230 |
|  | AMPE | 0.643 | 0.436 | 0.000 |
|  | Response Onset Time | 3.767 | 0.073 | 0.147 |
|  | Peak Speed | 5.640 | **0.032** | 0.225 |
|  | Peak Y-Force | 4.185 | 0.060 | 0.166 |
|  | Path Length | 7.950 | **0.014** | 0.303 |
| P2 | Duration | 3.721 | 0.074 | 0.145 |
|  | AMPE | 0.021 | 0.886 | 0.000 |
|  | Response Onset Time | 4.601 | 0.050 | 0.184 |
|  | Peak Speed | 3.141 | 0.098 | 0.118 |
|  | Peak Y-Force | 2.929 | 0.109 | 0.108 |
|  | Path Length | 2.833 | 0.114 | 0.103 |

**Figure 7.**
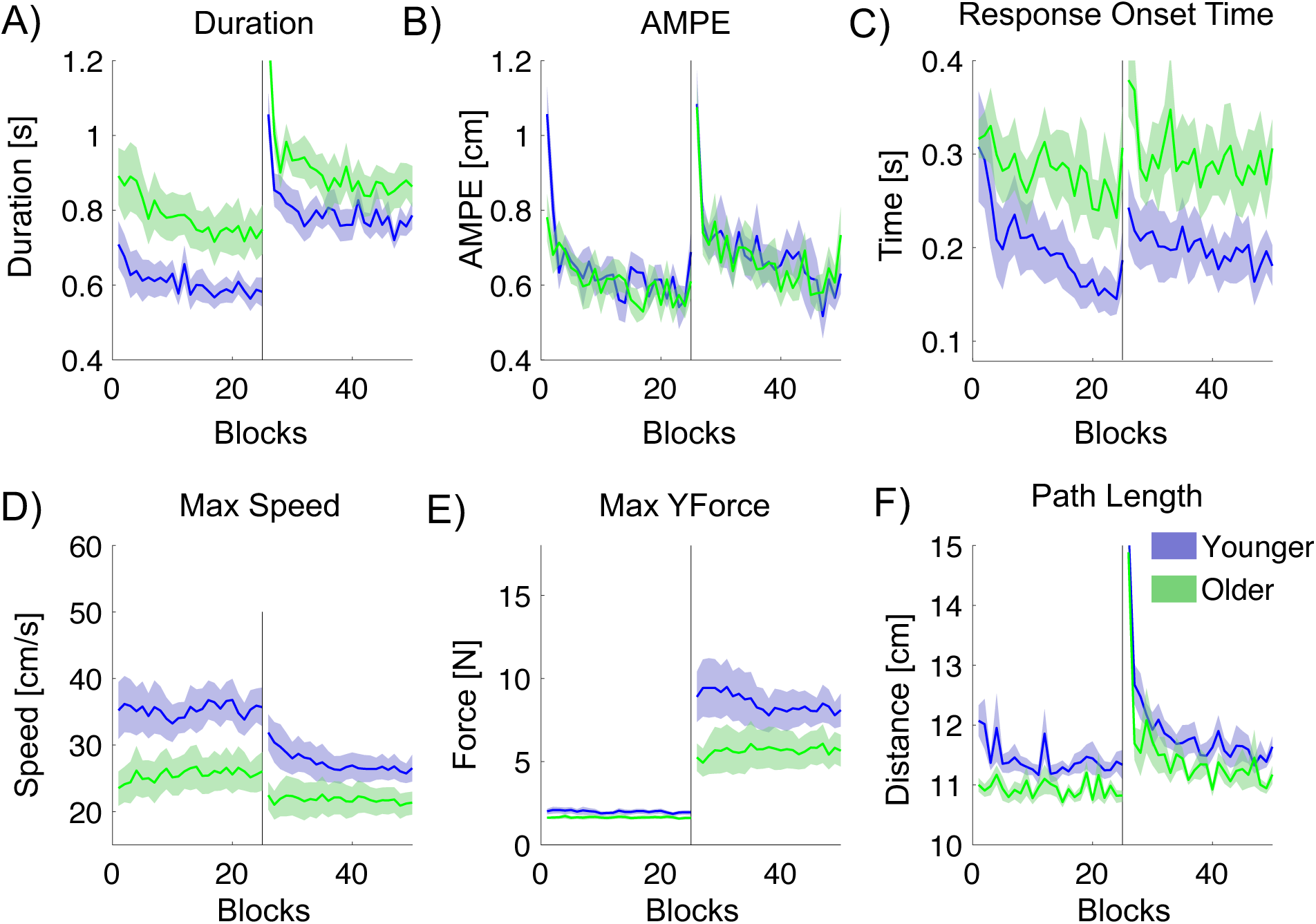
Movement metrics quantifying the hand trajectory during the mass center-out task, plotted against blocks. Younger participants are shown in blue, and older participants in green. The black vertical line indicates the midpoint of the experiment, at which we introduce a simulated 5 kg mass at the handle. **A:** Movement duration. **B:** Absolute maximum perpendicular error (AMPE). **C:** Response onset time. **D:** Maximum trajectory speed. **E:** Maximum force in the direction of movement (YForce). **F:** Trajectory path length.

### 4.3. Experiment 3: Handle Viscous Resistance Center-Out Movement - age related performance

**Figure 8.**
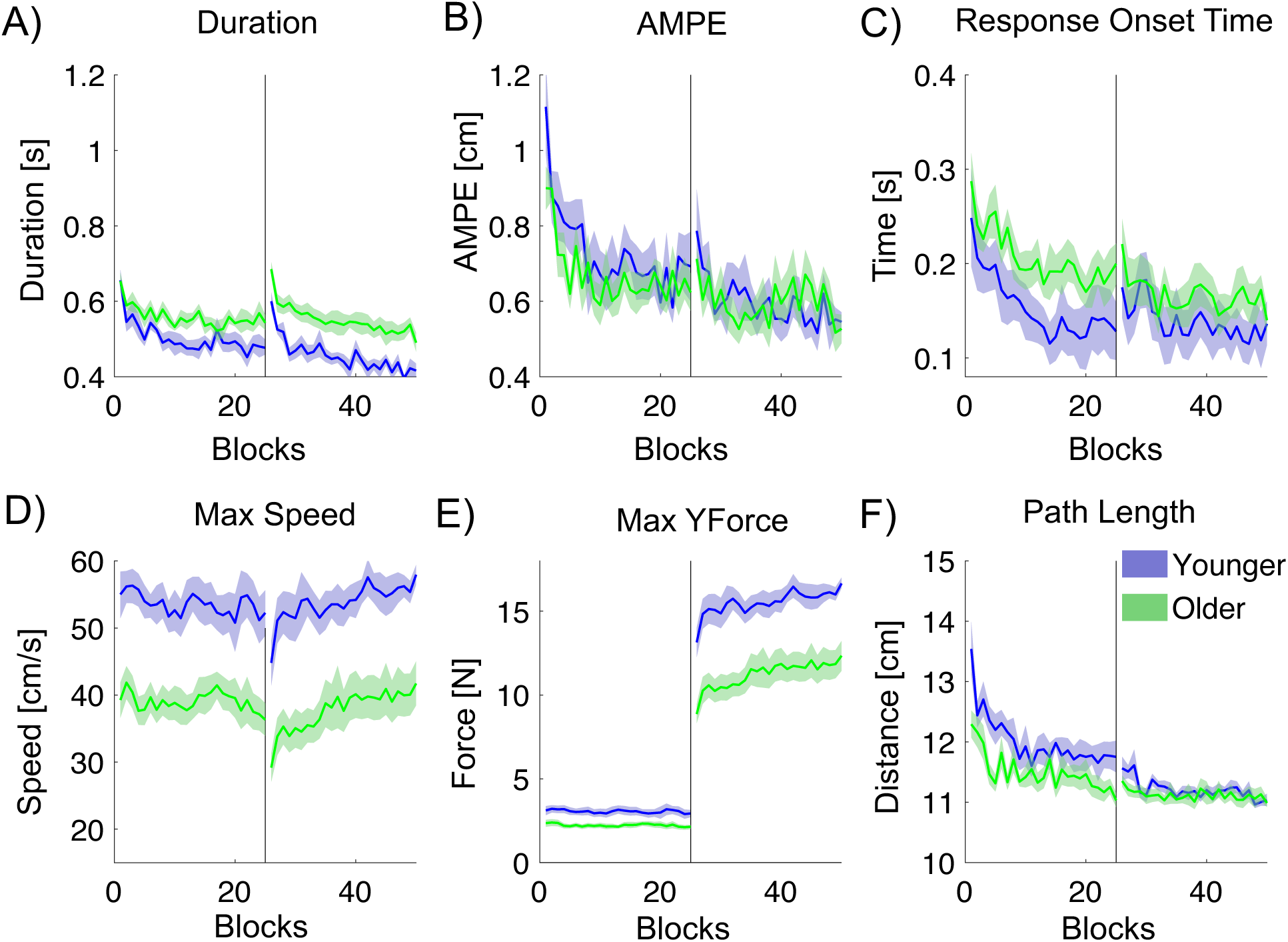
Movement metrics quantifying the hand trajectory during the right-hand viscous-field center-out task, plotted against blocks. Younger participants are shown in blue, and older participants in green. The black vertical line indicates the midpoint of the experiment, at which we introduce a simulated viscosity at the handle. A: Movement duration. B: Absolute maximum perpendicular error (AMPE). C: Response onset time. D: Maximum trajectory speed. E: Maximum force in the direction of movement. F: Trajectory path length.

Participants again performed a right-hand center-out task involving 12 cm movements from a central start position to one of eight targets, under two conditions: a null field and a viscous field. The task comprised two phases of 200 trials each to assess performance in unloaded and loaded movements.

Fig. 8 illustrates the performance of younger and older participants across multiple metrics that characterize hand trajectories during the null-viscosity center-out task.

Relative to the older group, younger participants complete the movements more quickly, exhibit shorter response onset times, reach higher peak speeds, apply greater force in the movement direction, and follow shorter paths. In contrast, AMPE remains comparable between the two groups

Repeated-measures ANOVAs were used to examine the significance of differences in the measured metrics between pairs of blocks (16 trials each) located at the initial null exposure, final null exposure, initial viscous dynamic exposure, and final viscous dynamic exposure. Post hoc comparisons were conducted to further assess significant differences between these phases.

Significant phase effects were observed across most metrics in both age groups. For younger participants, Duration, AMPE, Peak Y-Force, and Path Length showed consistent changes across all task phases. Response Onset Time and Peak Speed also changed significantly over time, although Response Onset Time was stable during the mass introduction phase. For older participants, strong effects were observed in Duration, AMPE, Peak Y-Force, and Path Length, with adaptation particularly evident between the null and viscous dynamic phases. Response Onset Time did not differ significantly across phases in this group. Notably, Peak Y-Force showed a large effect sizes in both groups (ω² > 0.85), indicating a strong sensitivity to the dynamic manipulation. Detailed results are summarized in Table 5.

For younger participants, Duration, AMPE, Response Onset Time, and Path Length decreased significantly during the null field phase. The introduction of the viscous dynamic led to significant increases in Duration, AMPE, Peak Y-Force, and Path Length, while Peak Speed decreased. By the end of the viscous phase, Duration, AMPE, Response Onset Time, and Path Length had decreased again, while Peak Speed and Peak Y-Force increased.

Younger participants showed more consistent adaptation across all measures, with significant changes at each phase, while older participants exhibited fewer significant changes, particularly in AMPE and Response Onset Time, and showed less sensitivity to the introduction of the viscous dynamic. This suggests younger adults adapted more readily to both the removal and addition of dynamic forces.

For older participants, Duration and AMPE decreased significantly during the null field phase. Duration increased with the viscous dynamic and then decreased again, while AMPE only decreased by the end. Peak Speed decreased with the dynamic and increased later, and Peak Y-Force increased throughout. Path Length decreased over both phases. Response Onset Time showed no significant changes.

**Table 5.** Repeated-measures ANOVA results for each age group with post hoc comparisons: ANOVAs were conducted for each performance metric within each age group across three key phases of the task. Bonferroni-corrected post hoc comparisons are reported for: (1) initial vs final null blocks, (2) final null blocks vs introduction of the viscous dynamic, and (3) start vs end of the viscous dynamic blocks. Significant p-values are shown in bold.

| AGE GROUP | METRIC | F | P | $\omega^2$ | P<sub>BONF</sub> (1) | P<sub>BONF</sub> (2) | P<sub>BONF</sub> (3) |
| --- | --- | --- | --- | --- | --- | --- | --- |
| Younger | Duration | 48.947 | < **0.001** | 0.463 | < **0.001** | < **0.001** | < **0.001** |
|  | AMPE | 50.677 | < **0.001** | 0.510 | < **0.001** | **0.001** | **0.002** |
|  | Response Onset Time | 18.853 | < **0.001** | 0.201 | **0.005** | 0.987 | **0.034** |
|  | Peak Speed | 18.361 | < **0.001** | 0.096 | 0.098 | 0.002 | < **0.001** |
|  | Peak Y-Force | 250.107 | < **0.001** | 0.855 | 0.059 | < 0.001 | < **0.001** |
|  | Path Length | 37.046 | < **0.001** | 0.555 | < **0.001** | < 0.001 | < **0.001** |
| Older | Duration | 53.759 | < **0.001** | 0.684 | < **0.001** | 0.141 | < **0.001** |
|  | AMPE | 31.797 | < **0.001** | 0.550 | < **0.001** | 0.141 | **0.027** |
|  | Response Onset Time | 4.953 | **0.005** | 0.063 | 0.253 | 0.868 | 0.236 |
|  | Peak Speed | 11.089 | < **0.001** | 0.116 | 1.000 | < **0.001** | < **0.001** |
|  | Peak Y-Force | 388.621 | < **0.001** | 0.916 | **1.000** | < **0.001** | < **0.001** |
|  | Path Length | 17.334 | < **0.001** | 0.401 | **0.009** | 1.000 | **0.007** |

### Across Age Groups Statistical Analyses

To investigate age-related differences in performance, we compared average values for each metric between younger and older participants using an ANOVA, analysing the first half of trials (1–200) and the second half (201–400) separately.

In Phase 1, significant age-related differences were found for Duration, Peak Speed, Peak Y-Force, and Path Length, with moderate to large effect sizes. In Phase 2, strong group differences were observed in Duration, Peak Speed, and Peak Y-Force, with large effect sizes (ω² > 0.47). No significant differences were observed for AMPE, Response Onset Time, or Path Length during this phase. Detailed results are summarized in Table 6.

**Table 6.** ANOVA results comparing younger and older participants within each experimental phase: Separate one-way ANOVAs were conducted for each performance metric in Phase 1 (P1, trials 1–200) and Phase 2 (P2, trials 201–400). Reported values include F-statistics, p-values, and effect sizes (ω²). Significant p-values are shown in bold.

| PHASE OF EXPERIMENT | METRIC | F | P | $\omega^2$ |
| --- | --- | --- | --- | --- |
| P1 | Duration | 6.411 | **0.025** | 0.265 |
|  | AMPE | 0.940 | 0.350 | 0.000 |
|  | Response Onset Time | 3.224 | 0.096 | 0.129 |
|  | Peak Speed | 16.316 | **0.001** | 0.505 |
|  | Peak Y-Force | 9.588 | **0.009** | 0.364 |
|  | Path Length | 4.703 | **0.049** | 0.198 |
| P2 | Duration | 14.343 | **0.002** | 0.471 |
|  | AMPE | 0.0004 | 0.984 | 0.000 |
|  | Response Onset Time | 0.844 | 0.375 | 0.000 |
|  | Peak Speed | 17.613 | **0.001** | 0.526 |
|  | Peak Y-Force | 18.594 | **< 0.001** | 0.540 |
|  | *Path Length* | 0.688 | 0.422 | 0.000 |

These qualitative results partially align with statistical analyses across the two age groups in the two phases, P1 and P2, of the experiments. Movement duration is significantly shorter in the younger participants (*p* = 0.025 and *p* = 0.002), who also exhibit higher peak movement speed, (p = 0.001 and p = 0.001) and greater force in the direction of motion, (*p* = 0.009 and p < 0.001). Path length is also significantly different in P1 (p = 0.049) No other metrics showed a statistically significant difference between the two age groups.

### 4.4. Experiment 3: Handle Viscous Resistance Center-Out Movement - Comparison Between Hands

**Figure 9.**
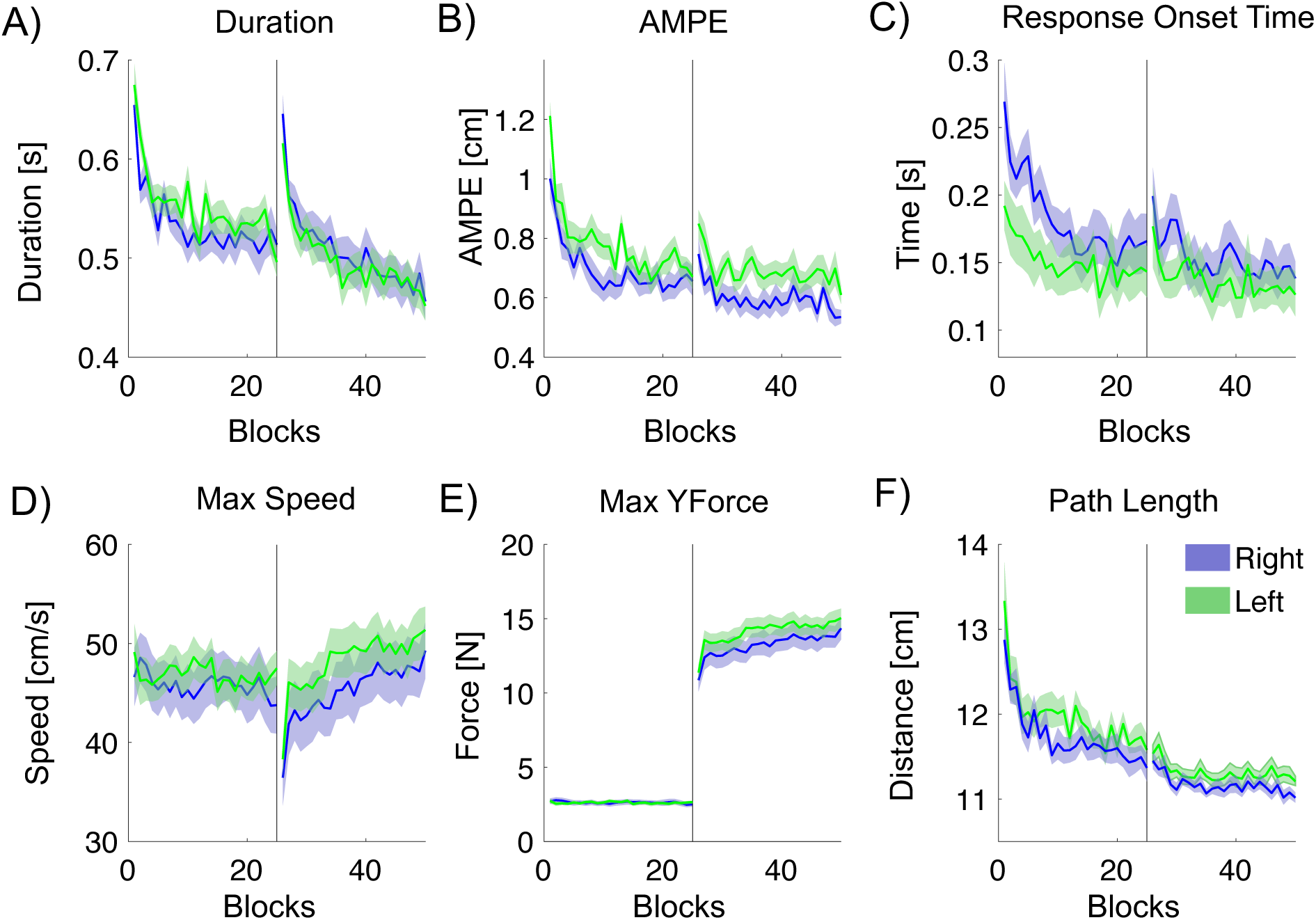
Movement metrics quantifying the hand trajectories during the null-viscous center-out task, plotted across blocks. The results from the experiments with participants using their right hand are shown in blue, and the corresponding results for the left-hand experiment are shown in green. The black vertical line indicates the midpoint of the experiment, at which we introduce a simulated viscosity at the handle. A: Movement duration. B: Absolute maximum perpendicular error (AMPE). C: Response Onset Time. D: Maximum trajectory speed. E: Maximum force in the direction of movement. F: Trajectory path length.

The experiment was then repeated with the left hand and a cross-hand comparison made. Fig. 9 shows the metrics across experimental trials, plotted as the mean ± SE. The most marked difference is that right-hand AMPE is notably lower than left-hand AMPE, although there were also other differences. Left-hand peak speed was higher than right-hand peak speed, and left-hand path length was longer. Statistical analyses confirm these observations.

To investigate differences in performance between the right and left hands, we compared average values for each metric between metrics using a repeated measures ANOVA, analysing the first half of trials (1 – 200) and the second half (201 – 400) separately.

In Phase 1, significant handedness-related differences were observed in AMPE, Response Onset Time, and Path Length, with AMPE showing the largest effect (ω² = 0.133). In Phase 2, AMPE, Response Onset Time, and Path Length also reached significance, though with smaller effect sizes. No significant differences were found in Peak Speed or Peak Y-Force during either phase. Detailed results are summarized in Table 7.

Right-hand AMPE values were significantly lower than left-hand values (*p* = 0.007 and p=0.01) and path lengths were significantly shorter (*p* = 0.046 and p = 0.016). No other metrics showed a statistically significant difference between the two hands. Response onset times were significantly shorter with the left hand (p = 0.002 and p = 0.042).

We note that the order of testing was fixed such that all participants performed the task first with their dominant arm, followed by the non-dominant arm. This design choice was motivated by the primary aim of the study, which was to examine age-related effects on dominant-arm control while minimising confounds in the mass condition. We acknowledge that this fixed order may have introduced order effects, as participants were more familiar with the task during the second session. However, despite prior task exposure, performance with the non-dominant arm was typically less accurate and more variable, suggesting that practice effects did not overcome inherent limitations associated with non-dominant limb control.

Interestingly, as shown in Fig. 9, movements performed with the right hand exhibited longer reaction times and lower movement speeds than those performed with the left hand. Because the non-dominant arm was tested second, biomechanical fatigue is unlikely to explain this pattern, although some degree of cognitive fatigue (e.g., reduced concentration) cannot be ruled out.

Given the relatively long experimental duration (30 minutes), one potential concern is the influence of fatigue on task performance. Prolonged task engagement could, in principle, reduce movement speed and force output, increase movement variability, and thereby impair movement accuracy. However, we did not observe any systematic degradation in performance over time in any of the measured metrics. On the contrary, performance generally improved across the experiment, consistent with ongoing task adaptation or learning. These observations suggest that fatigue was not a dominant factor influencing the results reported here.

**Table 7.** ANOVA results comparing younger and older participants within each experimental phase (third dataset): One-way ANOVAs were conducted separately for Phase 1 (P1, trials 1–200) and Phase 2 (P2, trials 201–400). The table reports F-values, p-values, and effect sizes (ω²). Significant p-values are shown in bold.

| PHASE OF EXPERIMENT | METRIC | F | P | $\omega^2$ |
| --- | --- | --- | --- | --- |
| P1 | Duration | 3.021 | 0.106 | 0.056 |
|  | AMPE | 10.167 | **0.007** | 0.133 |
|  | Response Onset Time | 14.011 | **0.002** | 0.047 |
|  | Peak Speed | 0.059 | 0.811 | 0.000 |
|  | Peak Y-Force | 0.143 | 0.712 | 0.000 |
|  | Path Length | 4.853 | **0.046** | 0.036 |
| P2 | Duration | 0.004 | 0.949 | 0.000 |
|  | AMPE | 9.138 | **0.010** | 0.151 |
|  | Response Onset Time | 5.104 | **0.042** | 0.012 |
|  | Peak Speed | 2.115 | 0.170 | 0.005 |
|  | Peak Y-Force | 3.514 | 0.083 | 0.010 |
|  | Path Length | 7.618 | **0.016** | 0.069 |

## 5. Discussion

### 5.1. The Effects of Aging on Sensorimotor Performance

In this study, we examined quantitative measures to test motor performance relevant to everyday tasks. We designed three experimental tasks using the vBOT robotic manipulandum (Howard et al., 2009), to accurately compare performance between healthy young adults (<35) and older adults (>60) in a controlled and repeatable fashion.

Across all three experiments, younger participants consistently demonstrated robust learning and adaptation in motor performance. They showed significant reductions in movement duration, Absolute Maximum Perpendicular Error (AMPE), Response Onset Time, and Path Length over the course of training, particularly in kinematic and viscous dynamic tasks. Younger individuals also adapted more consistently and quickly to the introduction of novel dynamics, such as added mass or viscous resistance, showing clear changes across all measured metrics. In comparison to older participants, younger adults had significantly shorter durations, higher peak speeds, greater directional force generation, and more extensive trajectory adjustments during the initial phases of exposure. These findings highlight a greater sensitivity to task dynamics and a higher capacity for motor adaptation among the younger cohort.

Analysis of right versus left-hand performance in Experiment 3 revealed significant asymmetries. AMPE and path length were both significantly lower for the right hand, indicating more accurate and efficient movements. In contrast, response onset times were significantly shorter with the left hand, possibly reflecting differences in motor planning or strategy rather than execution. No significant differences were found in peak speed or peak force between the hands. These hand-specific performance differences underscore the sensitivity and relevance of our methodology in capturing fine-grained variations in motor behavior. The ability to detect these subtle asymmetries demonstrates the utility of our approach for quantifying motor performance with high resolution and for identifying individual differences in motor control strategies.

The three experimental tasks differed significantly in their capacity to reveal age-related sensorimotor differences. Experiment 1, involving novel bimanual control of a virtual arm, highlighted initial difficulties among older adults in acquiring new motor skills, though performance improved significantly with practice. Experiment 2 assessed the management of inertial loads, clearly demonstrating older adults’ reductions in peak forces and prolonged movement durations. However, Experiment 3, featuring viscous resistance and a scoring system designed to reward brisk movements, was particularly effective at differentiating younger from older adults. By explicitly encouraging participants to move quickly, the scoring mechanism appeared to push individuals closer to their limits, thereby likely exaggerating performance differences. Consequently, this task identified pronounced age-related deficits in force production, response onset time, and movement duration, suggesting its superior effectiveness for clinical assessment of subtle motor impairments associated with aging.

The results presented here align with findings of previous studies utilizing robot tasks. For instance, Herter et al. demonstrated that robotic tools can detect subtle age-related declines in proprioception, reaction time, and motor coordination (Herter et al., 2014). Our findings that movements take longer and slow down with age are consistent with results by (Moulton et al., 2022). In addition, Kitchen et al. report that older adults show slower movement speeds, reduced peak forces, and impaired adaptation when exposed to perturbations using robotic systems (Kitchen and Miall, 2019), which strongly mirrors our findings.

Individual variability was found to be high, and some older adults performed similarly to younger adults. Variability may result from differences in lifestyle, regular physical activity, cognitive engagement, or other unexplored genetic and environmental factors (Hunter et al., 2016). As individuals age, sensorimotor performance often declines (Hunter et al., 2016; Yoshimura et al., 2020), impacting everyday activities such as drinking a glass of water or walking down the street (Seidler et al., 2010). This decline is largely due to age-related alterations in the brain and body that affect motor control (Clark and Taylor, 2011).

Across the three experiments, our findings align with multiple theoretical accounts of neuromuscular aging. First, age-related slowing in force production is predicted to result in increased movement durations (Hunter et al., 2016); consistent with this, older adults showed significantly longer movement times across all tasks, particularly in the viscous-field conditions where higher force generation was required. Second, theories of reduced peak muscular output with age predict lower maximum forces during dynamic movements (Power et al., 2013); this was confirmed in Experiment 3, where older adults consistently produced lower peak forces than younger adults. Third, neuromuscular decline is associated with a reduced capacity to adjust motor commands when task dynamics change (Seidler et al., 2010). Our results support this prediction: when transitioning from the null-field to the mass or viscous environments, older adults exhibited smaller improvements and less pronounced changes in movement speed, response onset time, and force generation, indicating reduced adaptability. Finally, theoretical models suggest that aging should impair de novo learning of novel motor mappings (Seidler et al., 2010); in Experiment 1, older adults demonstrated slower acquisition of the bimanual control strategy, consistent with this account. Together, these findings provide empirical confirmation of several hallmark features of neuromuscular aging.

The decline in sensorimotor performance with aging is thought to arise from multifaceted neural mechanisms, primarily involving structural and functional changes within the central nervous system. One of the most significant factors contributing to sensorimotor decline is a reduction in neuroplasticity - the brain’s ability to adapt and reorganize in response to new experiences or environmental changes (Burke and Barnes, 2006; Kourosh-Arami et al., 2021; Marzola et al., 2023). Additionally, brain volume tends to decrease over time (Raz et al., 1998; Fjell and Walhovd, 2010), which can impair the regulation of motor regions, further contributing to motor dysfunction (Seidler et al., 2010).

One of the prominent neural changes is the progressive loss of gray matter volume, particularly affecting regions such as the motor cortex, basal ganglia, and cerebellum, which are critical for motor planning, execution, and coordination (Seidler et al., 2010). Moreover, aging leads to reduced integrity of white matter tracts, including corticospinal and sensorimotor pathways, resulting in impaired signal conduction, reduced cortical excitability, and increased variability in motor responses (Cassady et al., 2020). A reduction in neurotransmitter availability - particularly dopamine - further impairs motor control, reduces precision, and limits the adaptability of motor responses, thereby contributing significantly to sensorimotor deficits in older adults (Kuehn et al., 2017).

In addition to neural deterioration, biomechanical factors substantially exacerbate age-related sensorimotor decline. Aging is accompanied by progressive muscle atrophy (sarcopenia), characterized by reductions in muscle mass, strength, and contractile speed, which significantly impair motor function and reaction times (Hunter et al., 2016). There is also a decline in proprioceptive acuity due to decreased density and sensitivity of peripheral sensory receptors (including muscle spindles and mechanoreceptors), impairing the ability to accurately sense joint position and movement—a capability crucial for precise motor control (Goble et al., 2011). Furthermore, increased joint stiffness and reduced flexibility, resulting from degenerative changes in connective tissues, limit movement range and fluidity, compounding sensorimotor impairment in older populations (Kulmala et al., 2014).

When motor function becomes further impaired by neurological diseases such as Parkinson’s disease, stroke, or other movement disorders, abnormal movements such as tremors (Deuschl et al., 1998), bradykinesia (Berardelli et al., 2001), rigidity (Fearon et al., 2015), or dyskinesia (Jankovic, 2005) become prevalent. These pathological conditions exaggerate the sensorimotor deficits already seen in healthy aging, introducing additional complexity for both assessment and rehabilitation (Dukelow et al., 2010; Edwards et al., 2012).

### 5.2. Current Clinical Paradigms

To assess human motor performance, several clinical paradigms have been developed. The clinical assessment “Timed Up & Go” (TUG) was designed as a brief evaluation of basic mobility skills in frail and older adults (Podsiadlo and Richardson, 1991). During the test, participants are instructed to stand up, walk three meters to a chair, and then sit down again. This assessment has proven to be an effective tool for evaluating mobility, walking, and balance (Shumway-Cook et al., 2000; Ng and Hui-Chan, 2005; Kristensen et al., 2007). Moreover, it has been demonstrated to be a reliable method for identifying adults at increased risk of falls (Nocera et al., 2013).

For the upper body, there has been continuous research aimed at developing tests that evaluate multiple aspects of motor function. Researchers have specifically focused on creating clinical assessments for upper-limb motor abilities. To evaluate upper-limb motor function, researchers introduced the Wolf Motor Function Test (WMFT) (Wolf et al., 2001), in which participants perform tasks that simulate daily activities, such as lifting or picking up objects. This test is widely used in hospitals, particularly for neurorehabilitation purposes (Fritz et al., 2009; Edwards et al., 2012; Turtle et al., 2020). Other well-known assessments include the Action Research Arm Test (ARAT), which evaluates post-stroke patients by measuring grasp, grip, pinch, and other motor tasks (Lyle, 1981). Another commonly used assessment is the Jebsen–Taylor Hand Function Test (JTHFT), also designed for post-stroke patients. This test is sensitive to changes over time, making it an effective tool for monitoring recovery (Jebsen et al., 1969).

Some tests assess both upper- and lower-limb movement. One example is the Fugl–Meyer Assessment (FMA), which is typically used to evaluate individuals with stroke or neurological impairments. It assesses balance and coordination in both the upper and lower extremities and is widely used in research and clinical settings (Sullivan et al., 2011).

### 5.3. Robotic Manipulanda Assessment

The experimental results presented here demonstrate that the vBOT system, in conjunction with the experimental procedures, can detect subtle changes in performance caused by age.

Previous research employing robotic manipulanda to assess sensorimotor ability has predominantly used standardized upper-limb reaching tasks designed to objectively evaluate specific aspects of motor and sensory function. Common movement paradigms include:

Center-out reaching tasks: Participants typically initiate movements from a central starting position to peripheral targets arranged radially around the starting point. These tasks assess motor accuracy, response onset time, movement speed, trajectory smoothness, and force control (Krebs et al., 1998; Bosecker et al., 2010).

Visuomotor adaptation paradigms: Participants perform reaching movements under altered visual feedback (e.g., rotated cursor) or novel force environments (e.g., viscous, inertial, or perturbation fields). Such tasks evaluate sensory integration, adaptation, learning rates, and the ability to generalize learned sensorimotor mappings (Shadmehr and Mussa-Ivaldi, 1994; Scott, 1999; Cressman and Henriques, 2009).

Bimanual coordination tasks: Tasks requiring simultaneous or coordinated movements of both arms have been used to investigate interlimb coordination, sensorimotor integration, and neural control mechanisms underlying complex motor behavior (Diedrichsen, 2007; Howard et al., 2008).

Proprioceptive assessment paradigms: Robotic manipulanda have been used to passively move a participant’s limb to specific positions, after which participants are asked to reproduce or discriminate limb positions. These paradigms quantify proprioceptive acuity, position sense, and kinesthetic deficits in both neurological disorders and aging populations (Dukelow et al., 2010; Herter et al., 2014).

Force generation and control tasks: Participants are asked to exert specified forces or match force profiles, either statically or dynamically. These tasks measure maximum voluntary force, accuracy of force modulation, and motor deficits related to muscle weakness or impaired neuromuscular control (Coderre et al., 2010).

Collectively, these paradigms enable detailed, quantitative assessments of various components of sensorimotor function, providing insights into motor control deficits, adaptation processes, and sensory impairments associated with aging, neurological disease, or injury. However, in cases of severe movement disruption—where patients may have difficulty initiating movement or are unable to perform certain tasks—additional or alternative metrics may be necessary to accurately assess functional capacity and impairment.

Compared to widely-used clinical assessments such as WMFT or ARAT, the robotic approach provides finer resolution and greater sensitivity, especially in detecting subtle deficits or tracking small changes over short intervals Clinically, the proposed robotic tasks could be implemented in outpatient rehabilitation settings, allowing practitioners to routinely quantify subtle sensorimotor deficits, monitor progression, and evaluate treatment efficacy. Such assessments could also be conducted using simpler devices, rather than the laboratory-based and costly vBOT. In particular, we are currently developing a low-cost robotic manipulandum for such purposes, based on an initial prototype (Howard, 2023). The target use case of such a device would be to provide a useful tool for clinicians and physiotherapists involved in patient rehabilitation, as well as for self-use for the patients themselves.

Emerging methods using robotic manipulanda, as utilized in this study, offer highly sensitive, quantitative assessments capable of detecting subtle impairments and characterizing abnormal movement patterns (Krebs et al., 1998). Future developments in robotic technologies combined with machine learning algorithms could further enhance diagnostic accuracy, enable precise tracking of disease progression, and allow personalized rehabilitation interventions tailored to the individual’s specific sensorimotor impairments (Coderre et al., 2010).

The three tasks used in this study were chosen to probe complementary aspects of sensorimotor control that are known to change with age. Experiment 1 assessed de novo bimanual learning of a novel kinematic mapping, demonstrating that although older adults showed slower movements and longer response onset times, they retained a clear capacity to learn the task. Experiments 2 and 3 examined adaptations to altered dynamics during familiar reaching movements, allowing age-related changes in force production, movement timing, and adaptability to be isolated more directly. The introduction of an inertial load in Experiment 2 revealed greater sensitivity in older adults to increased mechanical demands, whereas the viscous resistance task in Experiment 3, combined with performance incentives, proved most sensitive to age-related differences across multiple metrics, including response onset time, force generation, movement speed, and path length. Together, these findings indicate that tasks involving velocity-dependent resistance and increased performance demands are particularly effective at revealing subtle age-related differences in sensorimotor control, while de novo learning tasks primarily capture preserved learning capacity with reduced efficiency in older adults.

### 5.4. Conclusions

This study demonstrates that robotic manipulanda, specifically the vBOT (Howard et al., 2009) can be used to sensitively and quantitatively assess age-related differences in sensorimotor control. Across three experimental tasks, older adults consistently exhibited deficits in movement timing, force generation, and response onset time, although some individual variability was observed. The use of both unimanual and bimanual paradigms enabled detailed characterization of performance, including lateralized differences between dominant and non-dominant limbs.

Our results reinforce the growing consensus that robotic systems can complement traditional clinical assessments by providing objective, high-resolution data on motor function. Such tools offer considerable promise for early detection of motor decline, tracking disease progression, and the development of rehabilitation interventions.

We note that our proof-of-concept study involved participants who were physically fit, particularly among the older participant group. This represents a sample skewed towards higher-functioning individuals. As a result, the observed differences between younger and older adults were probably smaller than those that would be seen in less able individuals, who are more reticent to volunteer for studies such as the one presented here.

The SE shading on the metric plots indicates considerable variability within both younger and older participant groups, with some older individuals performing at levels comparable to younger adults. This variability highlights substantial individual differences in age-related sensorimotor performance and illustrates the limitations of purely cross-sectional comparisons. Longitudinal studies following the same participants over time would provide a clearer characterization of aging-related changes, although such approaches are not always practical. Nevertheless, acknowledging this variability is important, as it supports the potential value of personalized intervention strategies.

In the current study, participant characteristics were reported in terms of mean age and standard deviation. Additional information such as height, weight, BMI, medical history, and physical activity levels would have provided further context for interpreting motor performance. In future studies, we will seek to include a more comprehensive set of participant characteristics, particularly measures related to physical activity. Obtaining more representative participant samples will be a priority in future research.

Further studies should also include more diverse participant groups and explore how robotic assessments can be integrated into clinical workflows for both age-related and neurological conditions. Degenerative neurological conditions such as cerebellar ataxia and Parkinson’s disease cause upper-limb movement impairments that substantially reduce quality of life. Quantitative kinematic measures, including workspace coverage and movement variability, may capture condition-specific deficits (e.g., reduced workspace after stroke, coordination errors in cerebellar disease, and bradykinesia in Parkinson’s disease) and reveal subtle abnormalities in the early stages of neurodegeneration. In future work, robotic manipulanda could be used to implement movement paradigms, such as viscous curl-field adaptation tasks, as outcome measures and potential markers of early diagnosis and long-term recovery, as well as to deliver tailored rehabilitation therapies directly.

## Declarations

Ethics approval and consent to participate

All participants were naïve to the objectives of the experiments and provided written informed consent prior to participation. All experiments followed the ethics protocol approved by the University of Plymouth’s Faculty Research Ethics and Integrity Committee.

## Consent for publication

All authors have approved the manuscript for submission.

Availability of data and materials

Upon acceptance, all relevant raw data will be deposited in a publicly accessible online repository.

## Authors’ contributions

LAH, ISH, GS, and EE conceived the study. LAH and ISH implemented the study, and LAH conducted data collection. Data analysis was carried out by LAH and ISH. The initial draft of the manuscript was written by LAH. All authors - LAH, ISH, GS, and EE - reviewed and edited the manuscript.

## AI usage statement

During the preparation of this work, LAH and ISH utilized ChatGPT-5 for proofreading. After using this tool/service, the authors reviewed and edited the content as needed and take full responsibility for the content of the publication.

